# Kainate receptors are critical for permissivity to sustained, disorganized, and network-wide pathological activity in the epileptic dentate gyrus

**DOI:** 10.64898/2026.02.11.705397

**Authors:** Lucas Goirand-Lopez, Pascal Benquet, Mariam Alharrach, Bruno Delord, Valérie Crépel

**Affiliations:** INMED, INSERM, Aix Marseille University, France; University of Rennes, Inserm-U1099, LTSI, Rennes, France; Institut des Systèmes Intelligents et de Robotique (ISIR), Sorbonne University, CNRS, Paris, France

## Abstract

In temporal lobe epilepsy, the dentate gyrus (DG) undergoes extensive reorganization through recurrent mossy fiber (rMF) sprouting, involving both AMPA and kainate receptors (KARs) at granule cell-to-granule cell (GC-GC) recurrent synapses. While KARs are known to enhance neural excitability, their specific contribution to the dynamics of the epileptogenic GC-GC network remains poorly understood. Here, by building and assessing a DG network model endowed with KARs, we revealed that the slow excitatory postsynaptic potentials (EPSPs) they mediate, in synergy with persistent sodium currents, profoundly reshape epileptogenic dynamics by facilitating the conditions for pathological activity, through two complementary mechanisms. Regarding input processing, KARs extended the window for temporal integration in GCs, favoring the initiation of network aberrant activities in response to more dispersed input patterns. Concerning network interactions, KARs reduced the number of sprouted connections needed for the development of epileptiform discharges. In fact, the extension of the structural and functional parameter region driving epileptiform activity was effective both for their initiation and, critically, their sustained maintenance. Moreover, KARs drove a striking transition in network behaviour, from partially organized collective dynamics to a highly disordered regime. In their presence, massive firing propagated through the DG network, disrupting spatial and temporal spiking patterns. This resulted in increased activity dimensionality and entropy, together with reduced mutual information between neurons. Altogether, our results suggest that KARs are not merely amplifiers of network excitation but critical determinants that fundamentally reshape the dynamical landscape, rendering the sprouted DG network permissive to self-sustaining pathological activity.

## Introduction

Temporal lobe epilepsy (TLE) is recognized as the most common form of epilepsy in adults (Helmstaedter et al., 2025; Téllez-Zenteno and Hernández-Ronquillo, 2012). It is characterized by substantial pathological changes in the hippocampus, particularly through mechanisms of reactive plasticity (Téllez-Zenteno and Hernández-Ronquillo, 2012). This plasticity is prominently observed in the dentate gyrus, where surviving neurons form aberrant connections, leading to significant network reorganization (Buckmaster, 2012; Gabriel, 2004; Represa et al., 1989; Sutula et al., 1989). A critical feature of this reorganization is the pathological sprouting of recurrent mossy fibers (rMFs), which establish aberrant excitatory synapses between dentate granule cells (GCs: GC-GC synapses). Previous studies have demonstrated that synaptic transmission within this aberrant recurrent network is mediated by both AMPA receptors (AMPARs) and kainate receptors (KARs) (Artinian et al., 2015, 2011; Epsztein et al., 2005).

Among KARs, GluK2-containing subunits have been identified as critical in enhancing network excitability and facilitating seizure activity (Matsuda et al., 2016; Mulle and Crépel, 2021; Peret et al., 2014). Preclinical studies have underscored the therapeutic potential of targeting KARs, demonstrating that their blockade significantly reduces seizure frequency in TLE mouse models and suppresses epileptiform discharges in organotypic hippocampal slice cultures prepared from surgically resected tissue of drug-resistant TLE patients (Baudouin et al., 2024; Boileau et al., 2023; Peret et al., 2014). Recent observations using single-cell-resolution calcium imaging have further elucidated the role of KARs in promoting epileptic network activity, particularly by recruiting neurons into large coactive assemblies (Goirand-Lopez et al., 2023).

Despite these findings, a critical question remains: how does the interplay between the degree of sprouted GC-GC connectivity and the contribution of KARs at these synapses determine the threshold and the dynamics of epileptiform activity? Experimentally probing this structural-functional interplay poses significant challenges, as experimental tools to independently manipulate both the density of GC-GC synapses and their AMPAR/KAR composition in animal models are currently limited. To address these complexities, we adapted the Santhakumar computational model of the dentate gyrus, which provides a detailed simulation of this brain region (Morgan et al., 2007; Santhakumar et al., 2005). The Santhakumar model is a representation of the dentate gyrus that consists of a reduced network of 527 cells, including 500 GCs, 15 mossy cells (MCs), 6 basket cells (BCs), and 6 HIPP (hilar perforant path-associated) cells, reflecting the cellular proportions of the biological dentate gyrus. Each cell is modeled with an intricate multi-compartment architecture that includes both somatic and dendritic elements. The initial network structure includes AMPAR-mediated excitatory and GABAergic inhibitory synapses arranged in a biologically realistic ring topology that reflects the natural lamellar organization and the GC-GC connectivity. This framework enables a systematic exploration of how different levels of GC-GC synapses (ranging from 0% to 50%) affect pathological network dynamics. However, the original model did not account for KAR-mediated transmission at GC-GC synapses, an essential aspect of the epileptic network. In the present study, we extended this model by incorporating KAR-mediated synaptic responses at GC-GC synapses and their amplification by INaP, consistent with experimental observations (Artinian et al., 2011; Epsztein et al., 2010). Our findings show that KAR-mediated transmission at GC-GC synapses acts as a critical driver of dentate gyrus epileptiform activity by lowering both structural and functional activation thresholds. In doing so, KARs transform transient, locally confined responses into sustained, network-wide reverberating activity, characterized by high-dimensional, disorganized population dynamics. Together, these effects demonstrate that KARs are key orchestrators that render the sprouted network highly vulnerable by drastically increasing its permissivity to self-sustained pathological activity.

## Results

Our work aimed to link synaptic transmission to network dynamics to explain how KAR-mediated signaling shapes DG epileptiform activity in TLE, building on a previous model (Morgan et al., 2007; Santhakumar et al., 2005). First, we developed and validated a GC-GC synapse model integrating AMPAR- and KAR-mediated excitatory postsynaptic potentials (EPSPs) kinetics with INaP, constrained by experimental patch-clamp recordings, to capture EPSP amplitude, dynamics, and voltage-dependent amplification (Artinian et al., 2015, 2011; Epsztein et al., 2010). Next, we assessed how receptor composition governs temporal integration at GC-GC synapses and quantified its impact on GC spiking. Finally, we embedded these mechanisms in a biophysically detailed DG network to test how KAR inclusion at GC-GC synapses modulates both the threshold for DG excitability and the spatiotemporal landscape of epileptiform activity.

### A novel biophysical model with KARs at sprouted GC-GC synapses

First, we developed a biophysical model of GC-GC synapses that incorporated both AMPARs and KARs with distinct conductances. The model parameters were derived from experimental data from patch-clamp recordings (Epsztein et al., 2010). The model incorporated two key features of KARs relative to AMPARs: slower conductance deactivation kinetics and a lower maximal conductance, consistent with previous experimental observations. Postsynaptic response amplitudes were scaled appropriately to match experimental EPSPs (Artinian et al., 2011; Epsztein et al., 2010). We incorporated the persistent sodium current (INaP), to account for the voltage-dependent amplification of EPSPs in these synapses, using steady-state activation from a previous modeling study (Lee, 2007), with parameters adjusted to fit our experimental data (see Methods). Among them, the time constant of activation was fitted using a simplified model (see Methods) to account for the amplification of EPSPs by INaP at subthreshold membrane potentials (Vsubthreshold) relative to those at resting membrane potential (Vrest). The quality of this fitting procedure was confirmed by comparing EPSPs generated by the model (mEPSPs; see Methods) with experimental recordings. Both AMPAR-(black) and KAR-mediated (red) mEPSPs (Figure 1A1, 1B1) closely matched the experimentally recorded EPSPs (Figure 1A2, 1B2), whether synaptic current waveforms were injected (Figure 1A1, 1A2) or inputs were stimulated to activate synapses (Figure 1B1, 1B2). In particular, KAR-mediated EPSPs’ larger amplification at subthreshold potentials as well as their slower decay, compared to AMPAR-mediated EPSPs, were clearly reproduced in the model. Therefore, the model captured key features, including amplitude, dynamics, and voltage-dependent properties, of both AMPAR- and KAR-mediated EPSPs (Epsztein et al., 2010), validating the parameter choices.

**Figure 1.**
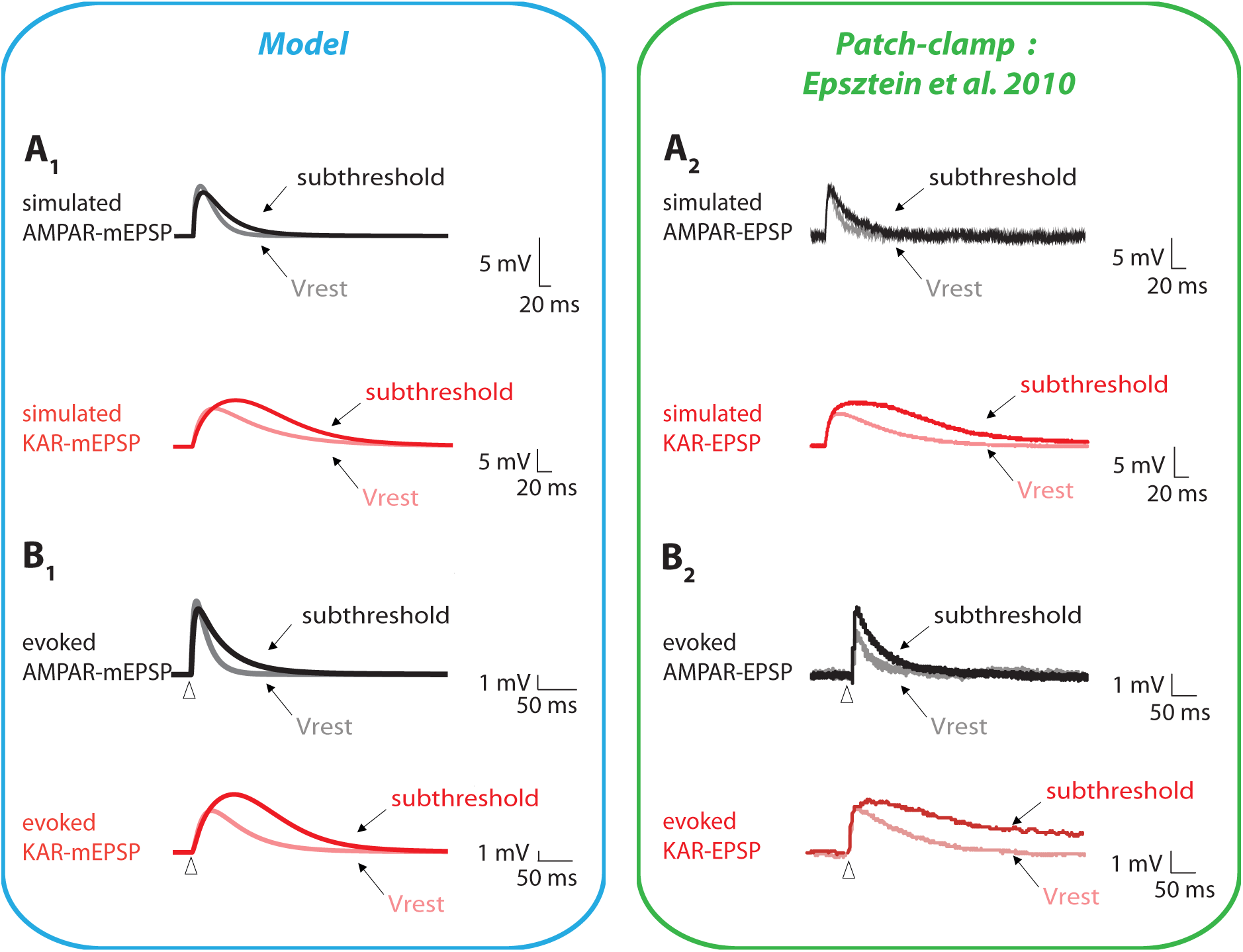
Experimental validation of model AMPAR- and KAR-mediated EPSPs. Each panel compares EPSPs under two receptor conditions: AMPAR-mediated (black) and KAR-mediated (red). Model (mEPSPs; A1, B1) and experimental (A2, B2) EPSPs were recorded either in conditions of resting (Vrest) or subthreshold membrane potentials and were offset-shifted for illustrative purposes. Simulated and evoked mEPSPs (A1, B1) accurately replicate their experimental EPSP counterparts (A2, B2), capturing both the distinct kinetics of the fast AMPAR-mediated EPSPs and slow KAR-mediated EPSPs, as well as the subthreshold amplification exhibited by the KAR-mediated EPSP compared to the AMPAR-mediated EPSP. In the model, simulated EPSPs were obtained by injecting a waveform mimicking the temporal envelope of AMPA and KAR synaptic currents (A1), while evoked EPSPs were obtained by activating AMPA and KAR synaptic gating dynamics (B1). In experimental patch-clamp recordings, simulated EPSPs were generated by a somatic current injection using a waveform (A2), while they were evoked by electrical stimulation positioned in the inner molecular layer of the dentate gyrus (B2).

We next assessed whether the model could account for the temporal integration of AMPAR- and KAR-mediated EPSPs and their impact on GC spiking, as observed experimentally (Artinian et al., 2011). Regarding temporal integration (Figure 2A1, 2A2), KAR-mediated mEPSPs (Figure 2A1, red) exhibited more pronounced summation in response to trains of synaptic stimulation at subthreshold membrane potentials, compared to AMPAR-mediated mEPSPs (Figure 2A1, black), as found experimentally (Figure 2A2) (Artinian et al., 2011). The model accounted for this effect across all conditions tested experimentally, i.e., at resting potential and subthreshold potentials, both with and without the INaP. Notably, the model reproduced the synergistic interaction between KARs and INaP, which significantly amplified EPSP summation at subthreshold potentials (Artinian et al., 2011).

**Figure 2.**
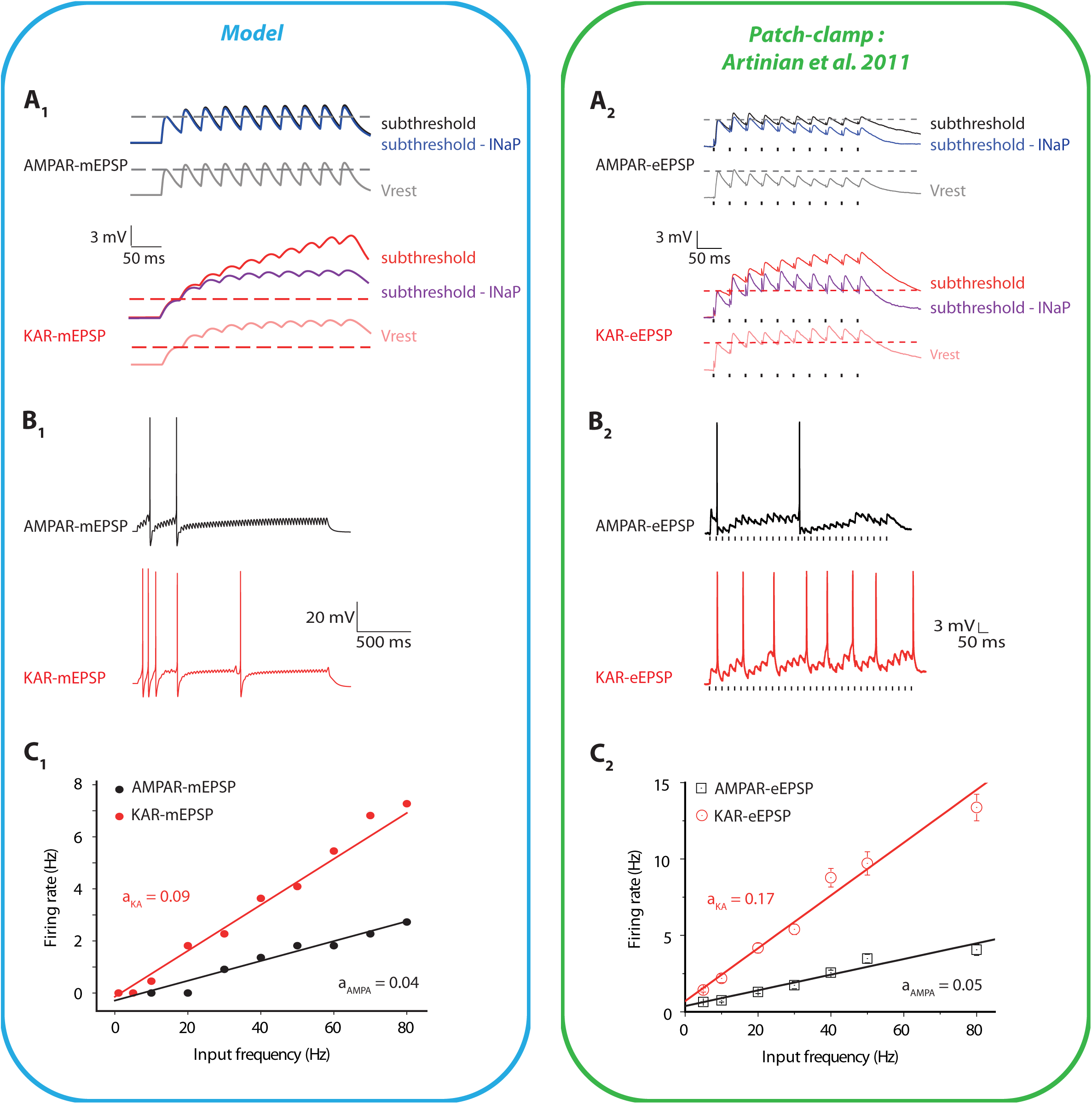
Experimental validation of model AMPAR- and KAR-mediated synaptic integration and firing dynamics. (A1-B2) Each panel compares EPSP integration upon 30 Hz input trains under either AMPAR-mediated or KAR-mediated conditions. (A1, A2) EPSP summation at resting (Vrest; AMPAR: light grey, KAR: light pink) and subthreshold potentials with (AMPAR: black, KAR: red) and without (AMPAR: dark blue, KAR: purple) the persistent sodium current (INaP). mEPSPs (A1) accurately replicate their experimental counterparts (A2), capturing the amplification exhibited by KAR-mediated EPSPs, relative to AMPAR-mediated EPSPs, upon subthreshold INaP activation. (B1, B2) Characteristic spiking dynamics upon 30 Hz input trains. The model (B1) replicates the differential firing patterns observed in patch-clamp recordings (B2). KAR-mediated mEPSPs (red) generate higher spike frequencies compared to AMPAR-mediated mEPSPs (black). (C1, C2) Input-output frequency relationships in the model (C1) and in patch-clamp recordings (C2). The model accounts for the increased input-output gain in response to KAR-mediated EPSPs, compared to the AMPA-mediated EPSPs.

We then examined the impact of the temporal integration of AMPAR- and KAR-mediated EPSPs on GC spiking in the model. mEPSP trains produced more spikes in the presence of KARs than in the presence of AMPARs (Figure 2B1), resulting in an input-output frequency relationship with a steeper slope (Figure 2C1), consistent with experimental data (Figure 2B2, 2C2) (Artinian et al., 2011).

Overall, the model thus accurately reproduced key effects of KARs, i.e., EPSPs with slower dynamics and synergistic amplification by INaP at subthreshold potentials, which, in turn, allowed greater temporal integration and higher output firing rates than those produced by AMPARs.

### KARs shift the spiking mode of model GCs from coincidence detection to integration over extended temporal windows

Building on the previous cellular-level validation, we designed an experimentally constrained, data-fitted GC model to explore how AMPARs and KARs influence network-level interactions. This framework enabled us to generate specific predictions about how these synaptic currents shape collective dynamics across the epileptic DG network. To investigate this, we first measured how AMPARs versus KARs affect GC spiking probability in the model sprouted network. We varied two parameters, the temporal window and in-degree (i.e., number of presynaptic GC inputs converging onto a single postsynaptic GC, Figure 3). For this analysis, we either considered AMPAR-only (*100%AMPAR* condition) or 50% AMPAR and 50% KAR (*50%KAR* condition), to match experimental conditions (Epsztein et al., 2005). We then examined how a single GC responded to inputs arriving within a defined temporal window of integration, mimicking phasic events. To capture the stochastic nature of neuronal activity, each presynaptic input was evoked at a random time within the temporal window. This modeling approach allowed us to independently manipulate two dimensions of the parameter region - namely, the width of the temporal window of integration and the number of GC-GC inputs - parameters that are not experimentally accessible.

**Figure 3.**
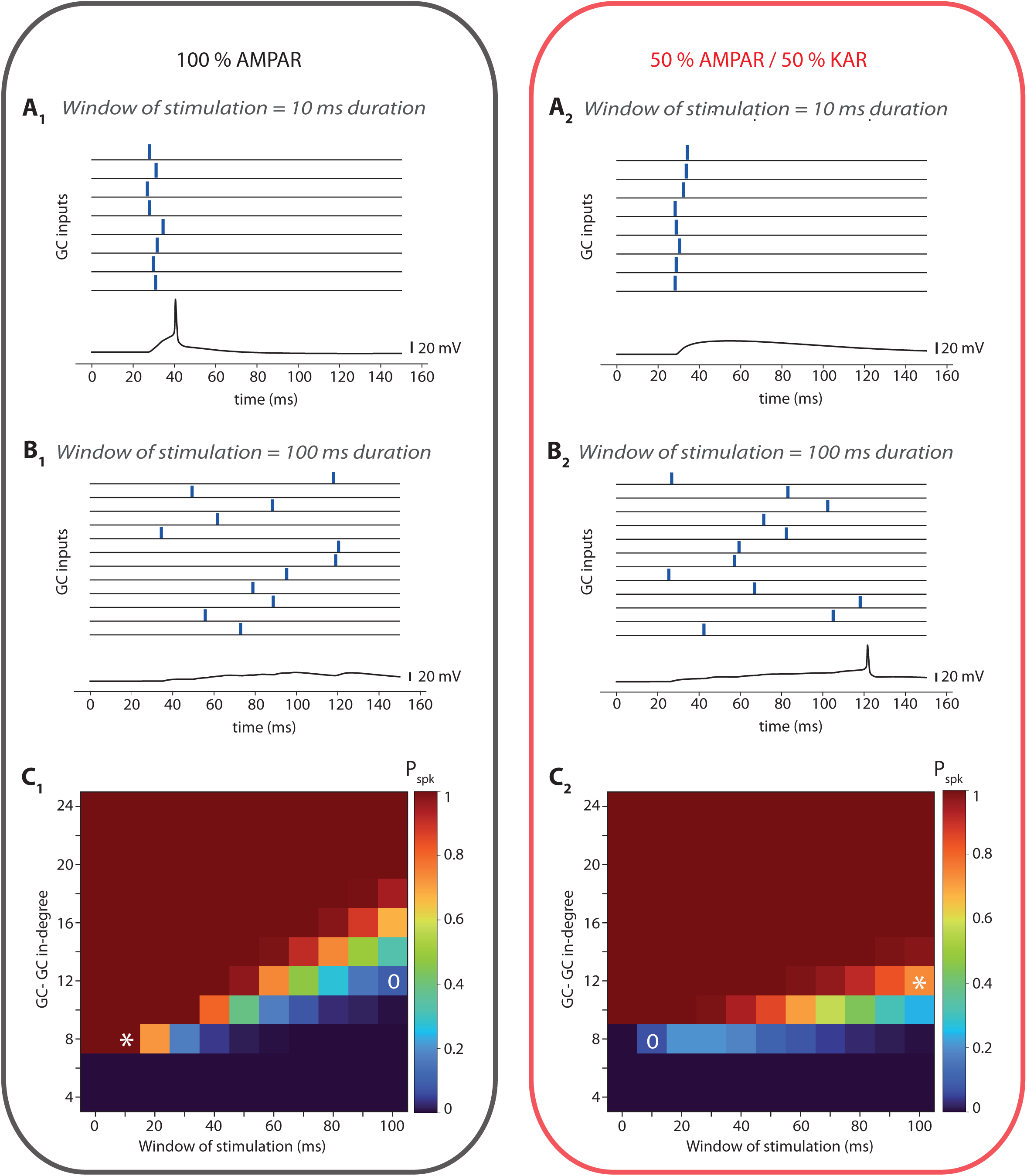
KARs extend the temporal integration windows for spike generation in response to inputs at sprouted GC-GC synapses. (A1-B2) Each panel compares EPSP summation and EPSP-spike coupling under either 100% AMPAR- or 50%AMPAR/50% KAR-mediated conditions. Top, example traces of GC inputs randomly delivered within a fixed temporal window: 8 inputs within 10 ms (A1, A2); 12 inputs within 100 ms (B1, B2). Bottom, corresponding GC responses. Under the AMPAR-only condition, a GC spike is triggered in response to the narrow input window (A1) but fails in response to the extended window (B1). Conversely, spiking occurs under the extended window (B2) but not the narrow one (A2) in the KAR condition. (C1, C2) Spike probability (Pspk) as a function of the number of inputs (GC-GC in-degree) and the duration of the input window, under AMPAR-only (C1) and mixed-receptor conditions (C2). Pspk values averaged over 1000 repetitions. In the AMPAR-only condition (C1), a small in-degree suffices to trigger spiking within narrow windows (around the star), while broader windows drastically reduce spike probability even at larger in-degrees (around the circle). Conversely, under mixed-receptor conditions (C2), narrow windows fail to trigger spiking at small in-degrees (around the circle), whereas broader windows enable reliable spiking at moderate in-degrees (around the star).

We observed that, at *100%AMPAR*, small temporal windows were sufficient for successful integration of inputs to generate postsynaptic GC spiking (Figure 3A1), whereas larger windows were not (Figure 3B1). In contrast, when synapses operated at *50% KAR*, spiking was promoted over broader temporal windows (Figure 3B2). Indeed, larger windows allowed sufficient time for the slow activation of INaP (Figure 2A), which was responsible for the sustained amplification and summation of successive incoming KAR-mediated EPSPs. At smaller windows, this mechanism was insufficient to compensate for the smaller amplitude of KAR-mediated EPSPs compared to AMPAR-mediated EPSPs (Figure 3A2).

To assess whether this trend was systematic, we computed the postsynaptic spike probability (Pspk) as a function of receptor type (AMPAR or AMPAR/KAR), window duration and number of presynaptic inputs (Figure 3C1, 3C2). This parametric analysis confirmed initial observations (Artinian et al., 2011). Indeed, for short windows (<30 ms), in the *100%AMPAR* condition, EPSPs (Figure 3C1) achieved higher spike probabilities than in the *50%KAR* condition (Figure 3C2). It also showed that synapses comprising only AMPARs required fewer inputs than those including KARs in triggering postsynaptic spikes. At longer windows (>30 ms), the mixed-receptor condition (*50%KAR*) significantly increased the probability of postsynaptic spikes (Figure 3C2) compared to AMPAR-operated synapses (Figure 3C1). Under these conditions, mixed-receptor-operated synapses required fewer converging inputs to reach the spike threshold compared with AMPAR-only synapses.

Globally, this parametric analysis revealed two key findings. First, AMPAR-endowed synapses enabled coincidence detection over short windows in GCs in response to perforant path (PP) inputs, in line with experimental observations (Leutgeb et al., 2007; Schmidt-Hieber et al., 2007). More specifically, the fast kinetics of AMPAR-mediated EPSPs conferred GCs with a high spiking propensity for narrow input windows (<30 ms), even at small in-degrees. Second, the incorporation of KARs extended the temporal window of integration, as found experimentally (Artinian et al., 2011; Epsztein et al., 2010). Notably, the model unravelled that KAR-endowed synapses enabled a more sensitive integration at longer timescales (>30 ms), i.e., requiring lower in-degrees to spike, compared to AMPAR synapses. Thus, the receptor composition at GC-GC synapses determines temporal-domain-specific sensitivity for spike generation.

### The sprouted DG network accounts for the diversity of epileptiform dynamics

We built a detailed biophysical model of the epileptic dentate gyrus (Figure 4A; see Methods and Supplementary file), using a 1:2000 scaling factor and comprising granule cells (GCs), basket cells (BCs), mossy cells (MCs), and HIPP cells (HCs), endowed with an extended set of ionic channels. Anatomically, the model was organized along three spatial dimensions. In the septo-temporal dimension (z-axis), the model exhibited a laminar organization (Figure 4B). In the two orthogonal dimensions (x- and y-axes), the model incorporated U-shaped lamellae of the dentate gyrus (DG) containing different neuronal types. This three-dimensional architecture enabled us to model local field potentials (mLFPs; Figure 4D-F), which arise from synaptic currents across all neuronal types at different laminar levels along the septo-temporal axis. Pathological rMF sprouting, i.e., the level of GC-GC out-degree connectivity, followed a small-world topology (Dyhrfjeld-Johnsen et al., 2007; Watts and Strogatz, 1998), with 5% random connections (Figure 4C).

**Figure 4.**
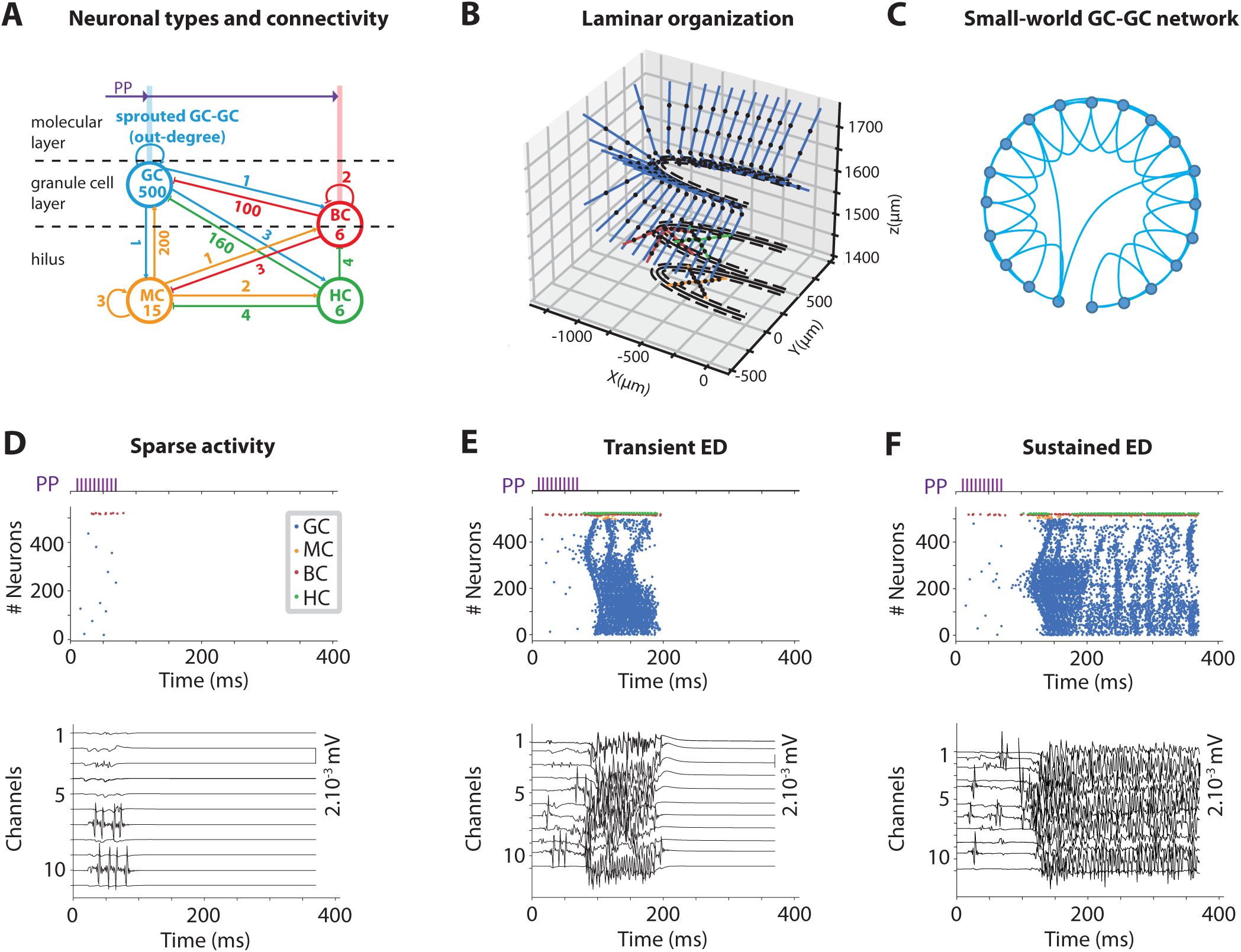
Structure, topology, and activity patterns in the dentate gyrus network model. (A) Schematic representation of the dentate gyrus (DG) network model. The model comprises four cell types distributed across three anatomical layers (molecular and granule cell layers and the hilus): granule cells (GC, blue), mossy cells (MC, orange), basket cells (BC, red) and hilar perforant path-associated cells (HC, green). The scheme indicates the cell numbers for each type, and the connectivity within and across layers, with their out-degree (number of output synapses per target neuron) indicated for each connection type. The number of sprouted GC-GC connections is a key variable parameter representing the extent of recurrent mossy fiber sprouting. (B) Three-dimensional representation of the model network, spatially organized in a laminar fashion along the septo-temporal dimension, and displaying the anatomical localization of all neuron types with representative examples of cells included in the computational DG model. (C) Schematic representation of the small-world GC-GC network topology of the epileptic DG. (D-F) Network responses to 10-phasic stimulations of perforant path (PP) inputs (purple), delivered within a 60 ms duration window at a sprouting level of 16%, and examined under AMPAR-only (D, E) or 50%AMPAR/50%KAR (F) conditions. Top, spike raster plots for the different cell types. Bottom, model local field potentials (mLFP) at channels along the septo-temporal dimension. The PP input could induce different activity patterns in the network model: (D) low-frequency, sparse responses; (E) transient responses with temporary widespread activation at higher firing frequencies (transient epileptiform discharge, ED); (F) sustained network responses with persistent activity across all cell populations (sustained ED), persisting for the duration of the whole simulation.

We monitored DG network dynamics in the model, in response to a train of 10 PP stimulations, delivered within different time windows to 1–2% of randomly selected GCs (Schneider et al., 2012) (see Methods). We assessed network activity by considering both GC spiking activity and mLFPs. When stimulated, the DG model could produce three distinct types of response (Figure 4D-F). First, with AMPAR-only, the model could display sparse spiking in a small subset of GCs and BCs that did not persist beyond PP stimulation (Figure 4D, top), indicating that this activity was induced by feedforward PP inputs and did not involve reverberation within the DG network. In this regime, BC firing followed the PP stimulation, while HCs and MCs remained essentially silent, consistent with insufficient recurrent excitation to engage the deeper inhibitory circuitry, as previously described (Santhakumar et al., 2005). This spiking activity resulted in mLFPs restricted to a small fraction of channels (Figure 4D, bottom).

The other two types of activity were characterized by the recruitment of a much larger fraction of GCs and BCs and the additional activation of MCs and HCs (Figure 4E, 4F, top). This spiking activity continued well after the termination of the PP stimulation, indicating that it was driven by network reverberation due to recurrent sprouted synapses. In this condition, mLFPs could be observed across all channels (see Figures 4E, 4F, bottom), with shapes resembling epileptiform discharges (EDs), as observed in an electroencephalogram (EEG) recording from an epileptic mouse (see Figure Supplement 1) and as previously reported (Dzhala and Staley, 2003; Peret et al., 2014). In ED regimes, BCs and HCs were robustly recruited alongside GCs and MCs, indicating that recurrent excitation was sufficient to engage the full inhibitory circuitry, again consistent with previous observations (Santhakumar et al., 2005). Remarkably, in these examples, the ED regime observed with AMPAR-only receptors rapidly vanished after PP stimulation, thereby qualifying it as transient ED (Figure 4E). In contrast, the third type of response, observed when KARs were included, consisted of reverberating, self-sustained network activity that persisted throughout the entire simulation, thereby characterizing it as sustained ED (Figure 4F). Altogether, these observations indicated that the architectural and biophysical constraints considered in the model allowed the emergence of the diversity of network dynamics, including distinctive epileptiform activities characterizing the hyperexcitable DG network.

### KARs broaden the parameter region driving epileptiform activity and enable sustained network hyperexcitability

In order to investigate how KARs influence the network’s propensity to generate epileptiform activity, we systematically assessed the probability of triggering EDs as a function of the parameters that set the timing of PP inputs (i.e., a fixed number of PP inputs across different window durations) and mossy fibre sprouting (i.e., GC-GC out-degree). We hypothesized that KAR-mediated transmission, with its distinct temporal integration properties (as demonstrated at the single-cell level, Figure 1-3), would affect the network’s ability to generate and maintain EDs. We assessed this hypothesis by analyzing the ED probability in the *100%AMPAR* (Figure 5A) and 50%KAR (Figure 5B) conditions.

**Figure 5.**
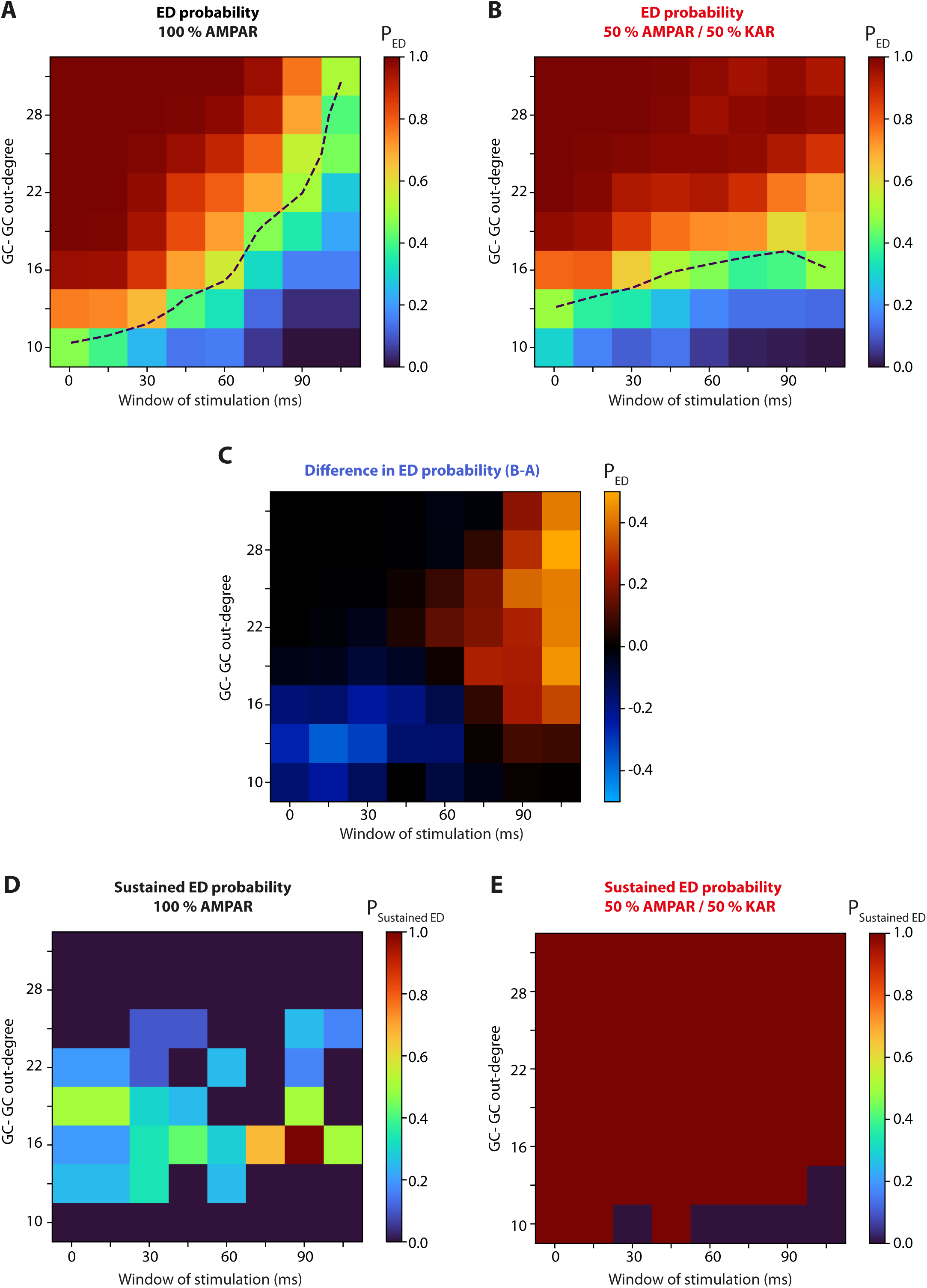
KARs facilitate the onset and propagation of epileptiform discharges over broad spatiotemporal regions. (A, B) Probability of generating EDs (P_ED_) under 100%AMPAR (A) and 50%AMPAR/50%KAR (B) conditions as a function of the duration of the stimulation window (see Fig. 4D-F, purple) and of the GC-GC sprouting level. P_ED_ values averaged over 10 repetitions. Black curves indicate the 50% ED probability isocline. (C) Difference in ED probability between the two conditions (i.e., (B)-(A)), highlighting regions in which KARs enhance (orange, positive values) or reduce (blue, negative values) P_ED_. (D, E) Probability of generating sustained EDs (P_sustained ED_) under 100%AMPAR (D) and 50%AMPAR/50%KAR (E) conditions.

In the *100%AMPAR* condition (Figure 5A), we observed that the ED probability was higher at short durations, i.e., for PP trains with higher frequency, inducing a higher level of AMPAR-mediated EPSP integration, due to their fast kinetics (see Figure 1-3). Consistently, longer windows impeded the integration of fast AMPAR-mediated EPSPs and decreased the probability of EDs. Also, increasing recurrent connectivity (GC-GC out-degrees > 10-15) raised ED probability. Moreover, both parameters affected ED probability to similar extents in the ranges considered (i.e., the slope of the 50% ED probability isocline, black dashed line) globally paralleled the first bisectrix). In the *50%KAR* condition (Figure 5B), we found that, contrary to the *100%AMPAR* condition, ED probability was less dependent on the duration of the stimulation window (as evidenced by a near-horizontal 50% isocline), consistent with the large time constant of KARs that allowed EPSP integration at lower frequencies, compared to AMPARs (see Figure 1-3).

To better visualize the specific effects of including 50% of KARs, we computed the difference between ED probability maps in their presence versus their absence (Figure 5C). The resulting representation revealed two parameter regions of interest: a large orange region indicating a higher ED probability when KARs were present and a small blue region of higher ED probability when only AMPARs were present. The small blue region indicated that triggering EDs in the *100%AMPAR* condition was favored for restricted windows (durations < 30 ms) and low sprouting level (GC-GC out-degrees ∼ 10-15). By contrast, the larger orange region highlighted that KARs shifted the region of higher probability towards longer windows (durations > 60 ms). Noticeably, in this condition, triggering EDs was strongly favored for a large range of sprouting levels, i.e., for all values of GC-GC out-degrees above 15. Overall, while at short window durations, the main factor favoring EDs was the sprouting level but not the nature of receptors (AMPARs vs KARs), our results revealed that, at longer stimulus durations, the major factor triggering EDs was the presence of KARs, independent of the sprouting level.

We then evaluated how KARs influence the temporal signature of EDs. We previously identified two distinct temporal signatures of epileptiform activities, i.e., transient (Figure 4E) and sustained (Figure 4F) EDs, which reflect increasing levels of activity reverberation in the DG epileptic network. We therefore systematically assessed the probability of sustained EDs (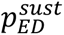; Figure 5D, 5E) as a function of the parameters setting the timing of PP inputs (window of stimulation) and network connectivity (GC-GC out-degree), in both *100%AMPAR* and *50%KAR* conditions.

In the *100%AMPAR* condition (Figure 5D), sustained EDs were globally very rare, being either absent or appearing at low probability 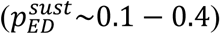 within a restricted region of the parameter space. Overall, this pattern suggests that under AMPAR-mediated transmission alone, sustained EDs occur only in a sparse manner across tested conditions. In striking contrast, the presence of KARs (Figure 5E) resulted in sustained EDs that occurred systematically 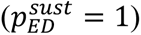 in the vast majority of the parameter region. Noticeably, even at lower sprouting levels (GC-GC out-degrees ∼ 10) and shorter stimulation windows (∼ 15-45 ms), the network could maintain sustained EDs, suggesting that the presence of KARs fundamentally enhances the network’s propensity to encounter lasting pathological activity, a marker of strong hyperexcitability.

Altogether, these findings highlight the crucial role of KARs in both the initiation (Figure 5A-C) and maintenance (Figure 5D, 5E) of epileptiform activity, consistent with previous single-cell (Figure 1-3) and network (Figure 4) results. The broader parameter region for ED generation and maintenance in KAR-endowed networks suggests that these receptors act as potent modulators of network excitability, markedly lowering the threshold for pathological synchronization.

### KARs drive a transition from structured to disordered network dynamics

To assess the spatiotemporal organization of sustained EDs in the presence or absence of KARs, we first visualized examples of spiking network activity dynamics (Figure 6A) as orbits in the principal component analysis (PCA) space of GC firing frequencies (Figure 6B). These orbits were colored as a function of the average topographical position encoded by the GC population’s activity. This representation revealed striking differences in collective dynamics between the two conditions. In the *100%AMPAR* condition, network dynamics orbited along an attractor exhibiting a large and well-structured shape (Figure 6B, left) that spanned an extensive range of encoded topographical positions (i.e., blue to red spectrum). Conversely, in the *50%KAR* condition (Figure 6B, right), dynamics revealed an unstructured attractor restricted to a much smaller region, together with a very limited range of encoded topographical positions (i.e., globally orange). Together, these observations indicated that KARs act as critical determinants of network organization. Indeed, whereas the sprouted network allowed organized activity and topographical encoding in the absence of KARs, their presence strongly disorganized activity and disrupted topographical encoding.

**Figure 6.**
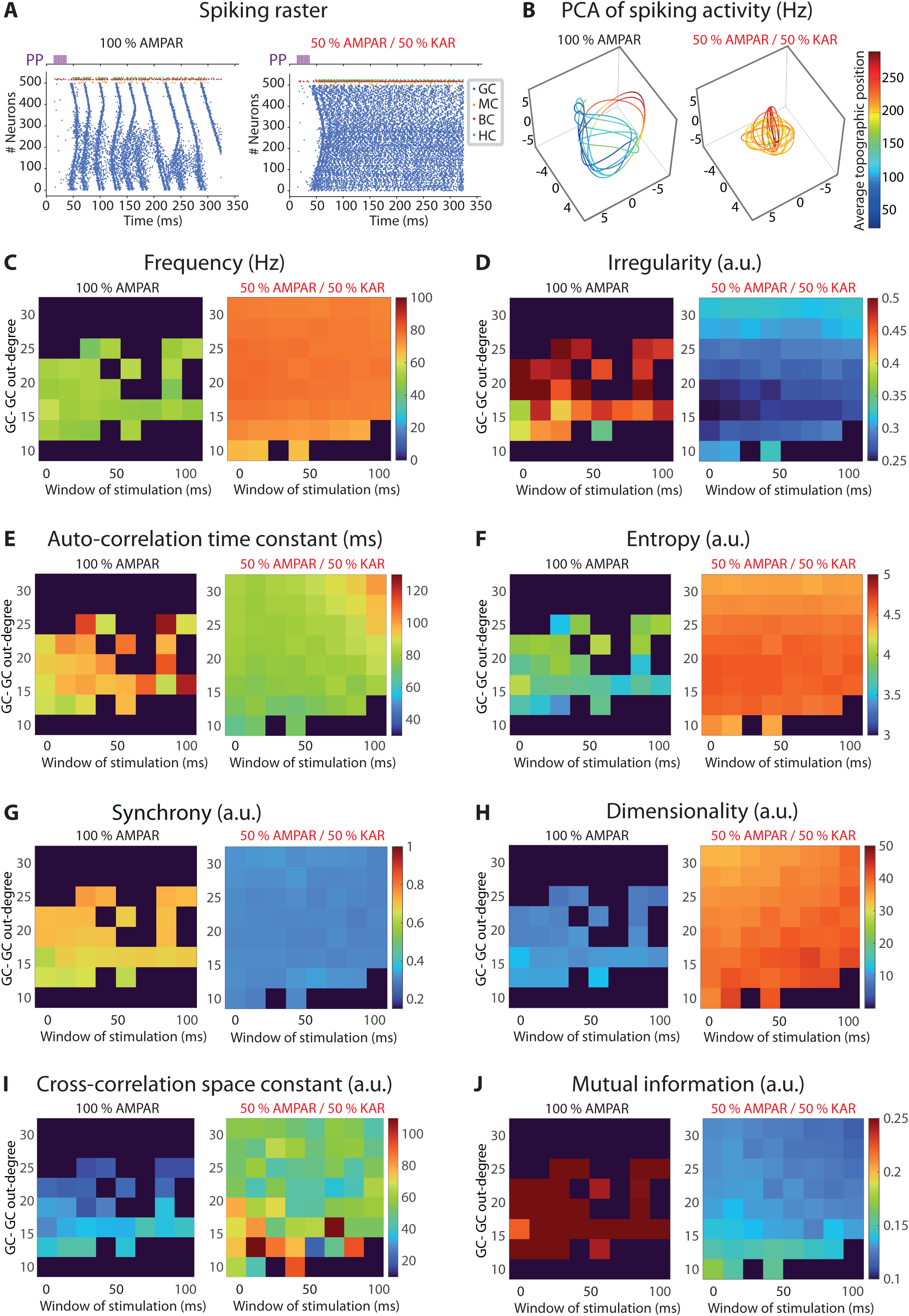
KARs shift DG dynamics from a partially organized to a highly disordered and complex regime. (A) Representative raster plots of the DG model network responses to a 10-phasic perforant path (PP) input stimulation (purple) delivered within a 30 ms duration window at a sprouting level of 25% under 100%AMPAR and 50%AMPAR/50%KAR conditions. (B) PCA visualization of firing rate trajectories corresponding to the rasters shown in (A). The color code represents the average topographical position of GC activity along the septo-temporal dimension. Neuronal (C-F) and network (G-J) level activity observables under both receptor conditions: neuronal mean firing frequency (C), spike irregularity (D), autocorrelation time constant (E) and entropy (F), and network synchrony (G), dimensionality (H), cross-correlation time constant (I), mutual information (J).

To better characterize DG dynamics in both conditions, we computed several observables to quantify GC activity at the single-neuron level (Figure 6C-F) and the network level (Figure 6G-J), as a function of stimulation window duration and sprouting degree. At the neuronal level, the presence of KARs led to a marked increase in spiking frequency (Figure 6C). Consistently, KARs induced a significant decrease in spiking irregularity (Figure 6D); this resulted from the degradation of the temporal organization of spiking, compared to the propagating bumps of activities observed in the *100%AMPAR* condition (Figure 6A), thereby reducing the temporal variability of spiking in individual neurons. Furthermore, the auto-correlation time constant of spiking was decreased in the presence of KARs (Figure 6E), revealing that this disorganization affected timescales up to the order of a hundred msec. These observations were further confirmed by an increase in entropy, reflecting greater unpredictability (Figure 6F).

At the level of network activity, we found that KARs strongly decreased the synchrony among neighbouring neurons (Figure 6G), indicative of a disorganization of the spatial structure of activity. Consistently, KARs drastically increased the dimensionality of collective spiking, i.e., more complex dynamics (Figure 6H). Moreover, the increase in the cross-correlation space constant in the presence of KARs (Figure 6I) further indicated a disorganization of activity at the spatial level, which arose from more global recurrent interactions across the network. Altogether, these changes in spatial organization of activity reduced the mutual information conveyed by collective interactions within the network (Figure 6J).

In summary, our simulations indicate that KARs profoundly transform ED network dynamics from a partially organized regime into a highly disordered and complex regime. Their presence in the DG network leads to a breakdown of temporal and spatial organization, generating high-dimensional pathological states with complex dynamics and severely disrupted coordinated activity.

## Discussion

The present modeling study unravels the mechanisms by which KARs contribute to epileptogenesis in the DG network, when endowed with aberrant sprouted GC-GC connections. We first reveal how the slower time-scale of KAR (vs. AMPAR) synaptic integration fundamentally lowers the threshold conditions for network epileptogenesis. Specifically, on the one hand, KARs render the DG network structurally permissive to aberrant activities, by requiring fewer sprouted GC-GC connections to drive epileptiform activities. On the other hand, KARs yield permissivity to aberrant activity in terms of responsiveness, by amplifying network reactivity to external inputs.

We then disclose that, whereas activity is restricted to local and transient spiking patterns under AMPAR transmission, KARs induce a state of collective spiking that invades the DG network both in the spatial and temporal domains, in the form of network-wide, sustained, pathological activity. Moreover, this massive propagation profoundly disorganizes collective spiking, accounting for a large repertoire of epileptiform activity properties, i.e., lowered temporal synchronization and increased dimensionality of network activity, leading to increased entropy levels and a collapse of mutual information. Altogether, our findings support the view that KARs do not induce a mere quantitative amplification of activity, but rather fundamentally shift the DG network into a different dynamical regime. Our results thus suggest that their presence constitutes a consubstantial determinant of the dynamical signature of the sprouted DG, rendering it more permissive and vulnerable to pathological self-sustaining activity. Therefore, KAR-mediated synaptic transmission appears as a critical lever for understanding epileptic DG network dysfunction.

### Novelty of the model

The present study aims to assess how GC-GC sprouted connectivity and the presence of KARs interact to set the threshold and dynamics of epileptiform activities. Probing this structure-function relationship experimentally poses significant challenges, as tools that allow independent manipulation of both the density of sprouted GC-GC synapses and their AMPAR/KAR composition are currently not available in animal models. In this context, modeling represents an essential means to address the question we study. Previous biophysical models (Morgan et al., 2007; Santhakumar et al., 2005) provided an account of DG activity by including AMPAR-mediated excitatory and GABAergic inhibitory synapses. However, they do not account for KAR-mediated transmission at sprouted GC-GC synapses, whereas it has been proven to be an essential feature of epileptic networks (Baudouin et al., 2024; Boileau et al., 2023; Matsuda et al., 2016; Peret et al., 2014). To address that limitation and to provide a more detailed account of DG activity, we leveraged previous studies to develop an original model incorporating, for the first time, KAR-mediated synaptic transmission at GC-GC sprouted synapses.

Moreover, we endowed the model with several additional novel properties, to further increase the biological relevance of our simulations. For instance, the anatomical description of the DG was improved, with the network being arranged in a multi-layered ring topology reflecting the natural lamellar organization of GC-GC connectivity, with dendrites being spatially oriented. Also, to establish a biophysically constrained implementation of recurrent interactions, we developed a detailed description of AMPAR- and KAR-mediated GC-GC synaptic transmission. Noticeably, the model captured two hallmark KAR properties, relative to AMPARs, i.e., slower decay kinetics and lower maximal conductance, and enabled the model to match experimentally observed EPSPs of both types. Additionally, GCs were endowed with the persistent sodium current (INaP), which preferentially amplifies KAR- (vs. AMPAR-) mediated synaptic transmission, in accordance with experimental observations (Artinian et al., 2015, 2011; Epsztein et al., 2010).

### Relevance and limitations of the model

The faithful reproduction of experimentally observed synaptic properties (Artinian et al., 2011; Epsztein et al., 2010) provided a critical validation and calibration step of the modeling approach, ensuring that subsequent cellular and network-level simulations were well grounded. The model was further validated by its ability to reproduce, at the network scale, the diversity of collective DG dynamics observed experimentally. Indeed, these mLFP responses emerged naturally, without any fitting procedure beyond that employed at the molecular scale of synaptic receptors, validating the biological realism of the multi-scale modeling approach. The model thus provides a computational framework for dissecting the complex interplay between network architecture, cellular biophysics, and emergent hyperexcitability in the epileptic DG, while offering a platform to test mechanistic hypotheses regarding the differential contributions of synaptic receptor subtypes to epileptogenesis.

Regarding possible future perspectives, the model itself could be improved by considering additional hypotheses, to capture the full complexity of molecular, cellular and network interactions at play in the epileptic DG. First, the scaling factor could be reduced to assess whether the present results still hold at the limit of larger orders of magnitude of the number of DG neurons. We expect that such upscaled versions of the model should display qualitatively similar regimes (although possibly shifted in the parameter space) in the presence or absence of KARs. Second, the model should be tested when incorporating the complete diversity of neuronal populations, instead of the four major cell types considered here, by adding, e.g., AAC (axo-axonic cells), MOPP (molecular layer interneurons with axons in perforant-path termination zone), HICAP (hilar interneurons with axons in the commissural/associational pathway termination zone) and IS (interneuron-selective cell types) (Dyhrfjeld-Johnsen et al., 2007; Morgan et al., 2007). Third, hilar neuronal death including mossy cells and somatostatin-positive interneurons (Buckmaster and Jongen-Rêlo, 1999; Ratzliff et al., 2002; Scharfman, 2019) could be assessed, as well as the incorporation of a small number of highly interconnected GC hubs (Morgan and Soltesz, 2008). For these structural manipulations, adding or deleting cells would involve small numbers of neurons compared to the total GC population. However, their contribution to global network dynamics has been shown to be substantial (Dyhrfjeld-Johnsen et al., 2007; Morgan and Soltesz, 2008). Nevertheless, the effects of KARs we unravel here are so massive that we expect our results to hold under such manipulations. Fourth, DG sclerosis is one of the main features observed in TLE (Blümcke et al., 2011; Mathern et al., 1996). Computational modeling of the epileptic DG reveals that sclerosis enhances small-world characteristics of the connectivity because of compensatory mossy fibre sprouting, which increases seizure susceptibility (Dyhrfjeld-Johnsen et al., 2007; Morgan and Soltesz, 2008). However, we anticipate that these structural changes would not affect our key findings on the role of KARs in the initiation and propagation of pathological DG activity, as KARs are expressed at recurrent GC-GC connections. Systematically testing the contribution of KARs to network hyperexcitability under these different pathological conditions (varying degrees of sclerosis, inclusion of specific hilar cell loss, and adding GC hub cells, etc.) opens up exciting avenues for future works.

### KARs transform GC-GC interactions in the sprouted DG

Our previous experimental work has shown that the slow kinetics of KAR-mediated EPSPs support extended temporal summation of synaptic inputs, whereas AMPAR-mediated EPSPs integrate over much narrower windows. Moreover, we previously reported that the recruitment of INaP amplifies the effective integration of EPSPs at larger time scales in the presence of KARs but not AMPARs. These effects resulted in an input-output frequency relationship with a steeper slope (Artinian et al., 2011). Our model was able to account for the effects of the interaction of KAR and INaP currents in broadening the effective integration window, enhancing the summation of successive inputs, and driving a larger neuronal gain.

We then assessed how these effects might manifest depending on the context in which the sprouted DG network operates. To accomplish this, we parametrically studied how GCs respond to irregular input patterns within a temporal integration window, thereby emulating the stochastic nature of neuronal spiking that signals phasic events. AMPAR-mediated transmission enabled coincidence detection when inputs were synchronized within short windows, consistent with the established role of GCs as precise temporal filters (Leutgeb et al., 2007; Schmidt-Hieber et al., 2007). By contrast, KARs weakened coincidence detection and enhanced temporal summation at longer time scales, by reducing the number of inputs required to reach threshold. Such transformation of integration properties likely contributes to the breakdown of the dentate gyrus’s normal “gatekeeper” function in the epileptic network (Hsu, 2007; Krook-Magnuson et al., 2015), wherein the typically sparse and selective firing of granule cells becomes hyperexcitable and less discriminative.

### KARs lower the threshold for DG epileptiform activity

At the network level, the model was able to successfully account for the diversity of DG spiking dynamics, including the simple sparse response to PP inputs, as well as transient or sustained EDs. These simulated ED patterns align with previous experimental observations of interictal activity exhibiting variable waveforms and durations (Dzhala and Staley, 2003). This showed that, under similar architectural constraints, the dynamical regime of the sprouted DG network was very sensitive to changes in initial conditions or biophysical model parameters. Our systematic exploration of the parameter space governing ED generation revealed that KARs fundamentally reshape the conditions under which epileptiform activity emerges and persists in the recurrent dentate gyrus network.

While AMPAR-only networks exhibited narrow parameter dependencies - requiring precisely timed, high-frequency inputs and moderate sprouting connectivity - the incorporation of KARs dramatically broadened the repertoire of conditions permissive for ED initiation, particularly at longer stimulation windows where slow KAR kinetics enable effective temporal summation. In line with this, KARs have been reported to promote spontaneous recurrent bursts in organotypic hippocampal slices exhibiting recurrent mossy fiber sprouting (Peret et al., 2014).

Most strikingly, KARs transformed the network’s propensity for sustained, hyperexcitable activity in the absence of any tonic external drive: whereas the AMPAR-mediated networks rarely maintained prolonged EDs across the parameter space, KAR-containing networks enabled sufficient levels of activity reverberation to maintain sustained epileptiform discharges under the vast majority of tested conditions, even at lower sprouting levels. Consistent with this result, KARs have been reported to prolong epileptiform events in cortical slices from naïve rats under pro-epileptogenic conditions (Szádeczky-Kardoss et al., 2019).

### KARs fundamentally set the dynamical regime of the sprouted DG network

When considering sustained EDs, our results demonstrate that KARs do not simply amplify excitation in the sprouted DG network, but rather shift network dynamics into a qualitatively different regime characterized by a profound degree of spatiotemporal disorganization. Indeed, at the single-neuron scale, KARs strongly disorganized the temporal structure of activity. This manifested as increased spiking regularity (reflecting a degradation of the temporal organization of spike trains), shortened autocorrelation time constants (revealing a disorganization at extended timescales), and increased entropy (indicating a greater unpredictability of discharge). These findings align with previous observations revealing that pathological hippocampal activities can emerge from reduced spike-timing reliability and decreased synchronization rather than from hypersynchrony (Foffani et al., 2007).

At the population level, KARs strongly disorganized the spatial structure of activity, reducing local synchrony, increasing the complexity of collective activity, broadening interactions and lowering conveyed mutual information across the network. In summary, our simulations indicated that KARs profoundly transform ED network dynamics from a partially organized pattern into a highly disordered and complex regime. Their presence in the DG network leads to a breakdown of the temporal and spatial organization, generating high-dimensional pathological states that severely disrupt activity coordination. Mechanistically, we propose that the slow kinetics of KARs sustain elevated firing through temporal summation, thereby stabilizing reverberation and sustained activity, while simultaneously disrupting the precise spike-timing coordination required for structured collective dynamics.

This dual KAR-mediated temporal and spatial reorganization of network dynamics is consistent with experimental observations showing that the transition from normal activity to seizure is characterized by a profound qualitative reconfiguration of collective dynamics, rather than a uniform increase in synchrony. The KAR-associated pattern we observe resonates with the canonical progression of focal epilepsy, in which seizure onset often begins with low-voltage fast activity that is relatively desynchronized before evolving into hypersynchronous activity (Bou Assi et al., 2020; Dzhala and Staley, 2003; Jiruska et al., 2013; Mormann et al., 2003; Wendling et al., 2003). Moreover, several studies report that dynamical complexity is elevated during interictal-like states yet decreases sharply at the ictal transition (Araújo et al., 2022; Jiruska et al., 2010), providing further evidence for qualitative shifts in network organization rather than monotonic changes in any single metric.

Taken together, these convergent lines of evidence suggest that KAR-mediated dynamics support a sustained, high-dimensional, desynchronized phase that precedes and facilitates the emergence of seizures. Within this framework, KARs would be pro-epileptogenic in three ways, by (i) lowering the threshold for epileptiform induction, (ii) stabilizing pathological reverberation, and (iii) shifting the network towards more complex, high-dimensional activity patterns, i.e., that could create a permissive substrate for seizures.

This interpretation is further supported by several studies demonstrating that selective blockade or reduction of GluK2-containing KARs robustly suppresses seizures in temporal lobe epilepsy models (Baudouin et al., 2024; Boileau et al., 2023; Matsuda et al., 2016; Peret et al., 2014).

## Materials and methods

The dentate gyrus network model we built was principally derived from previous modeling and experimental studies (Morgan et al., 2007; Santhakumar et al., 2005; Tejada et al., 2014, 2012; Tejada and Roque, 2014; Yim et al., 2015). Anatomically, it was spatially organized as U-shaped lamellae stacked along the septo-temporal dimension, which allowed estimating model local field potentials (mLFP). The network was endowed with four neuronal types, two of them being excitatory - granule cells (GCs) and hilar mossy cells (MCs) - and two inhibitory - basket cells (BCs) and HIPP cells (HCs) (Santhakumar et al., 2005). It was scaled with a 1:2000 ratio of the number of neurons, compared to the rodent dentate gyrus, with 500 GCs, 15 MCs, 6 BCs, and 6 HCs. Each cell type was described as a biophysically realistic multicompartmental model that included a soma and a dendritic tree (containing either 2 or 4 dendrites) and was endowed with multiple ionic and synaptic modeled conductances. The connections between different cell types were as previously described (Santhakumar et al., 2005). In this network architecture, each PP fibre contacted on average 1 GC and 1.2 BCs: PP-BC sparsity is 0.2 (Yim et al., 2015) and the model comprises 6 BCs. Network stimulation was achieved by applying a short train of 10 PP spikes, with a duration of up to 105 ms, to 5–10 randomly selected GCs (i.e., 1–2% of the 500 GCs), and 1 spike per PP over BCs on average. Synaptic weights were adjusted so that single presynaptic spikes reliably elicited postsynaptic spikes. The multicompartmental architecture, intrinsic and synaptic modeled properties, based on specific morphological and electrophysiological data of the four cell types reported in the literature, as well as the procedures for estimating mLFP and activity observables can be found in the Supplementary File. The model was simulated in the NEURON environment (Hines and Carnevale, 1997). Modeling data were computed via a custom-made MATLAB/Python code (see below).

## Acknowledgements

We thank Alfonso Represa for helpful comments and discussions.

## Additional information

### Author contributions

VC designed the original architecture of the project. VC and LGL designed the study of dentate gyrus cellular aspects. VC, BD and LGL designed the study of dentate gyrus network aspects. LGL built and simulated the model. PB and MA supervised and LGL achieved the study of model local field potentials. BD supervised, and BD and LGL achieved the analysis of the network activity. VC and BD wrote the manuscript with contributions from LGL and feedback from PB and MA.

### Data availability

Modeling data were generated via a custom-made MATLAB/Python code available through GitHub at https://github.com/lgoirandlopez/EpilepticDG3D500GCsLFPMeasure. The experimental data used in the present study were previously published in Epsztein et al. (2010) and Artinian et al. (2011) as indicated in the text and figures.

