## Supplemental methods and figure for "Kainate receptors are critical for permissivity to sustained, disorganized, and network-wide pathological activity in the epileptic dentate gyrus"

**Supplementary methods**

***Model of the dentate gyrus***

**Compartmental voltage dynamics**

Each neuron was described as multicompartmental, comprising a soma and 2 to 4 dendrites segmented into several compartments. Each compartment $j$ of the dendrite $k$ was modelled as an equivalent cylinder with diameter $d_{j,k}$ and length $L_{j,k}$. Its membrane potential, $V_{j,k}$, evolved according to an approximation of the cable equation (Hines and Carnevale, 1997), calculated at the axial midpoint of the compartment

$$c_{m,j,k}\frac{dV_{j,k}}{dt} + s_{j,k}(\sum_i I^{ions}_{i,j,k} + \sum_i I^{syn}_{i,j,k}) = \frac{V_{j-1,k}-V_{j,k}}{r_{j-1,j,k}} + \frac{V_{j+1,k}-V_{j,k}}{r_{j,j+1,k}},$$

where $V_{j-1}$, $V_j$ and $V_{j+1}$ are the membrane potential of successive compartments (along an axis from the soma to the tip of the considered dendrite $k$, one of the two terms on the right-hand side being absent at the soma or the tip of the dendrite), $\sum_i I_{ions,i,j,k}$ and $\sum_i I_{syn,\,i,j,k}$ the sum of the ionic and synaptic current densities – described below – flowing through the membrane of that compartment, and where

$$c_{m,j,k} = C_m s_{j,k}$$

is the membrane capacitance of compartment $j$ of dendrite $k$, with $C_m$ being the membrane specific capacitance,

$$s_{j,k} = \pi d_{j,k} L_{j,k}$$

the compartment surface area, and axial resistances between successive $x$ and $y$ compartments of dendrite $k$ defined as

$$r_{x,y,k} = \frac{2Ra}{\pi}\left(\frac{L_{x,k}}{d^2_{x,k}} + \frac{L_{y,k}}{d^2_{y,k}}\right),$$

with axial resistivity $Ra$.

In the case of somatic membrane potential, $V_{soma}$, there was no compartment before and several after (corresponding to the number of dendrites). The somatic membrane potential, $V_{soma}$ evolved following :

$$c_{m,soma}\frac{dV_{soma}}{dt} + s_{soma}(\sum_i I^{ions}_{i,soma} + \sum_i I^{syn}_{i,soma}) = \sum_k \frac{V_{1,k}-V_{soma}}{r_{1,k,soma}},$$

with conventions as described above.

**Ionic currents of neurons**

The voltage– or calcium–gated ionic current described below were modelled using the Hodgkin-Huxley formalism. Hence, their activation and inactivation gating variables $x$ evolved according to first-order kinetics

$$\frac{dx}{dt} = \frac{x_\infty - x}{\tau_x},$$

with their steady-state

$$x_\infty = \frac{\alpha_x}{\alpha_x+\beta_x},$$

and time constant

$$\tau_x = \frac{1}{Q_{10}(\alpha_x+\beta_x)},$$

depending on voltage- and/or calcium-dependent (i.e., depending on currents) gating kinetic coefficients setting the transition rates from the closed to the open state ($\alpha_x$) and from the open to the closed to state ($\beta_x$), and $Q_{10}$ being the temperature coefficient, following, unless stated,

$$Q_{10} = 3^{\frac{T-6.3-273.15}{10}},$$

with $T$ the température, expressed in Kelvin degrees. For most gating variables, the model was set by giving either kinetic coefficients $\alpha_x$ and $\beta_x$ or both the steady-state and time constant dependances, unless stated. In the following, $R = 8.3134\ J.mol^{-1}.K^{-1}$. $R = 8.3134\ J.mol^{-1}.K^{-1}$, $F = 96485\ C.mol^{-1}$. The exact composition of currents depended on the cell type and are listed below, after the definition of current models.

**Fast sodium current (NaT)**

The fast sodium current is responsible for the rapid depolarization of the action potential. Its model equations were adapted from previous works (Aradi and Holmes, 1999; Santhakumar et al., 2005; Tejada et al., 2014). The current followed

$$I_{NaT} = \overline{g_{NaT}} m^3 h\left(V - V_{Na}\right),$$

with $V_{Na}$ the reversal potential for sodium currents and $\overline{g_{NaT}}$ the maximal sodium conductance. Gating was defined by

$$\alpha_m(V) = \frac{0.3(V+43)}{1-\mathrm{exp}exp\left(-\frac{(V+43)}{5}\right)}$$

and

$$\beta_m(V) =- \frac{0.3(V+15)}{1-\mathrm{exp}exp\left(\frac{(V+15)}{5}\right)}$$

for the activation gate, and

$$\alpha_h(V) = \frac{0.23}{\mathrm{exp}exp\left(\frac{(V+65)}{20}\right)}$$

and

$$\beta_h(V) = \frac{3.33(V+15)}{1+\mathrm{exp}exp\left(-\frac{(V+12.5)}{10}\right)}$$

for the inactivation gate.

**Persistent sodium current (NaP)**

Previous work has shown that the persistent sodium current activates at subthreshold potentials during EPSPs (Artinian et al., 2011; Epsztein et al., 2010) mediated by kainate receptors. We incorporated persistent current into the model by adapting equations from previous models (Lee, 2007) to match the dynamics observed experimentally (Artinian et al., 2011; Epsztein et al., 2010). The NaP current followed

$$I_{NaP} = \overline{g_{NaP}} m_{NaP} h_{NaP}\left(V - V_{Na}\right)$$

Activation kinetics was defined by

$$m_{NaP\infty}(V) = \frac{1}{1+exp\left(-\frac{v+48.5}{3.4}\right)} ,$$

and

$$\tau_{mNaP}(V) = k_{NaP}\left(0.025 + 0.14exp\left(\frac{V+40}{10}\right)\right), \ if\ V <- 40mV$$

$$\tau_{mNaP}(V) = k_{NaP}\left(0.02 + 0.145exp\left(-\frac{V+40}{10}\right)\right), \ if\ V \geq -40mV ,$$

where $k_{NaP} = 525$ is a kinetic coefficient optimized (see procedure below). Inactivation was set by

$$h_{NaP\infty}(V) = \frac{1}{1+exp\left(\frac{v+15.5}{13.3}\right)},$$

and the function $\tau_{hNap}$ was estimated from published data[11,21]

$$\tau_{hNap}(V) = 2000 + 3920exp\left(-0.5\left(\frac{V+68}{29}^{2}\right)\right).$$

**Fast and slow delayed rectifier potassium currents (Kf and Ks)**

The fast and slow delayed rectifier potassium currents are involved in the repolarization of the membrane potential following an action potential. Both current models were adapted from previous works (Aradi and Holmes, 1999; Santhakumar et al., 2005; Tejada et al., 2014) and followed

$$I_{Kx} = \overline{g_{Kx}}x^{4}\left(V - V_{K}\right), \ x = \{f, s\},$$

with first-order gating kinetics and kinetic coefficients

$$\alpha_{f}(V) = \frac{0.07(V+18)}{1-exp(-\frac{(V+18)}{6})}$$

and

$$\beta_{f}(V) =- \frac{0.264}{exp(\frac{V+47}{40})}$$

for the fast delayed rectifier current and

$$\alpha_s(V) = \frac{0.028(V+30)}{1-exp(-\frac{(V+30)}{6})}$$

and

$$\beta_s(V) =- \frac{0.1056}{exp(\frac{V+55}{40})}$$

for the slow delayed rectifier current.

**A-type potassium current (A)**

The A-type potassium current is fast-activating, inactivating and is crucial for delaying the onset of action potential, attenuating the excitability and modulating the beginning of repolarization. Its model is adapted from previous works (Aradi and Holmes, 1999; Santhakumar et al., 2005; Tejada et al., 2014) and follows

$$I_A = \overline{g_A} kl(V - V_K),$$

with particular gating kinetics, with steady-states

$$x_\infty(V) = \frac{1}{1+\alpha_x(V)}, \quad x = \{k, l\}$$

and time constants

$$\tau_s = \frac{\beta_x}{Q_{10} a_{0x}(1+\alpha_x)}$$

with $a_{0k} = 0.08$ and $a_{0l} = 0.02$, and

$$\alpha_k(V) =- 3.10^{-3}(V + 33.6)\frac{9.648.10^4}{RT}$$

and

$$\beta_k(V) =- 1.8.10^{-3}(V + 33.6)\frac{9.648.10^4}{RT}$$

for the activation gate, and

$$\alpha_l(V) = \beta_l(V) = 4.10^{-3}(V + 83)\frac{9.648.10^4}{RT}.$$

**Voltage- and calcium-dependent big potassium current (BK)**

BK potassium currents are voltage- and calcium-dependent, with a high unitary conductance contributing to final spike repolarization and fast afterhyperpolarization. The current model was adapted from previous work (Tejada et al., 2014) and followed

$$I_{BK} = \overline{g_{BK}}o(V - V_K)$$

with gating set by

$$\alpha_o(V, [Ca^{2+}]) = 0.28[Ca^{2+}]/(\left[Ca^{2+}\right] + 0.48.10^{-3}exp\left(-\frac{2*0.84F}{RT}V\right))$$

$$\beta_o(V, [Ca^{2+}]) = 0.48/(1 + [Ca^{2+}]/(13.10^{-6}exp(-\frac{-2F}{RT}V)).$$

**Voltage- and calcium-dependent small potassium current (SK)**

SK currents are weakly voltage-dependent and highly calcium-dependent potassium, with a low unitary conductance. They contribute to the slow afterhyperpolarization and to spike-frequency adaptation. Their model was adapted from previous work (Yim et al., 2015) and followed

$$I_{SK} = \overline{g_{SK}}q(V, t)^2\left(V - V_K\right)$$

with a specific kinetic scheme with $\alpha_q\left(\left[Ca^{2+}\right]\right) = 0.00246\, exp(\frac{12log_{10}([Ca^{2+}])+28.48}{4.5})$

and

$$\beta_q\left(\left[Ca^{2+}\right]\right) = 0.006\, exp(-\frac{12log_{10}([Ca^{2+}])+60.4}{35})$$

**Inward rectifier potassium current (Kir)**

The inward rectifier current contributes to the membrane potential near the resting state. Epileptic GCs exhibited an upregulated inward-rectifier K+ current (Young et al., 2009). The current model was taken from published work (Yim et al., 2015) and followed

$$i_{Kir} = \overline{g_{Kir}}l\left(V - V_K\right)$$

with steady-state

$$l_{\infty}(V) = \frac{1}{1+exp\left(\frac{V+98.92}{10.89}\right)}$$

and time constant

$$\tau_{l}(V) = \frac{1}{Q_{10}\left(0.006exp\left(-\frac{-V}{67.08}\right)+0.08*exp\left(\frac{-V}{67.08}\right)\right)}$$

the $Q10$ temperature coefficient (ms) being in that case,

$$Q_{10} = 1^{(T-33-273.15)/10}.$$

**T–type calcium current**

The transient and low-threshold T–type calcium current notably contributes to the after depolarizing potential (ADP). We adapted its model from a previous model [6]. The current followed

$$I_{CaT} = \overline{g_{TCa}}a^2b\Delta V_{T}$$

with $\Delta V_{T}$ being a driving force following a Goldman-Hodgkin-Katz-like equation

$$\Delta V_{T} = f\left(1 - \frac{[Ca^{2+}]_{inside}}{[Ca^{2+}]_{outside}}exp(\frac{V}{f})\right)\frac{V}{f}\frac{1}{1-exp(\frac{V}{f})}$$

with

$$f = 10^{3}\frac{RT}{2F}$$

and gating variables dynamics set by

$$\alpha_{a}(V) = \frac{0.2(V-19.26)}{1-exp(-\frac{(V-19.26)}{10})}$$

and

$$\beta_{a}(V) = \frac{0.009}{exp(-\frac{V}{22.03})}$$

for the activation gate, and

$$\alpha_{b}(V) = 10^{6}exp(-\frac{V}{16.26})$$

and

$$\beta_b(V) = \frac{1}{1+exp(-\frac{V-29.79}{10})}$$

for the inactivation gate.

**N–type calcium channel**

The transient, high-threshold N–type calcium current contributes to large calcium spikes. Its model was adapted from a previous model (Tejada et al., 2014) and followed

$$I_{CaN} = \overline{g_{NCa}} c^2 d(V - V_{Ca})$$

with $V_{Ca}$ the reversal potential set by the Nernst Equation

$$V_{Ca} = 10^3 \frac{RT}{2F} log\left(\frac{[Ca^{2+}]_{outside}}{[Ca^{2+}]_{inside}}\right)$$

with $[Ca^{2+}]_{outside} = 2mM,$ with gating kinetic coefficients expressing as

$$\alpha_c(V) = \frac{0.19(V-19.88)}{1-exp(-\frac{(V-19.88)}{10})}$$

and

$$\beta_c(V) = 0.046exp(-\frac{V}{20.73})$$

for the activation gate, and

$$\alpha_d(V) = \frac{0.00016}{exp(-\frac{V}{48.4})}$$

and

$$\beta_d(V) = \frac{1}{1+exp(-\frac{V-30}{10})}$$

for the inactivation gate.

**L–type calcium channel**

The sustained L–type calcium current is activated by prolonged depolarization. It was taken from a previous model (Tejada et al., 2014) and followed

$$I_{CaL} = \overline{g_{CaL}} e^2 \frac{0.01}{0.01+[Ca^{2+}]} \Delta V_{CaL}$$

with the driving force following a Goldman-Hodgkin-Katz like equations :

$$\Delta V_{CaL} = f\left(1 - \frac{[Ca^{2+}]_{inside}}{[Ca^{2+}]_{outside}} exp(\frac{V}{f})\right)\frac{V}{f}\frac{1}{1-exp(\frac{V}{f})}$$

with $f$ defined as for the T–type calcium current and gating defined by

$$\alpha_e(V) = 15.69 \frac{(V-81.5)}{1-exp(-\frac{(V-81.5)}{10})}$$

and

$$\beta_e(V) = 0.29 exp(-\frac{V}{10.86}).$$

**Calcium dynamics ($Ca^{2+}$)**

Calcium dynamics resulted from calcium inflow through the T–, N– and L–type calcium channels, and a passive decay mimicking extrusion and/or buffering. The intracellular calcium concentration $[Ca^{2+}]_i$ thus evolved according to (Aradi and Holmes, 1999)

$$\frac{d[Ca^{2+}]}{dt} = \frac{10^7}{dF} \sum_{X=\{T,N,L\}} ICa_X - \frac{[Ca^{2+}]-[Ca^{2+}]_0}{\tau_{Ca}}$$

with $d = 200\, nm$ the thickness of the membrane, $10^7$ a conversion coefficient from $A.m^{-2}$ to $mA.cm^{-2}$, $\tau_{Ca} = 10\, ms$ and $[Ca^{2+}]_0 = 0.05\mu M.$ is the basal intracellular calcium concentration.

**Hyperpolarization activated fast and slow currents (Hf and Hs)**

The cationic fast and slow H currents are non-selective and activated by hyperpolarization. We adapted the two currents from a previous model (Tejada et al., 2014). The fast H current followed

$$I_{Hx} = \overline{g_{Hx}} h_f^{\,2}(V - V_{Hx}),\ \ x = \{f, s\}$$

with gating set by

$$h_{f\infty}(V) = \frac{1}{1+exp(\frac{V+91}{10})}$$

and

$$\tau_{h_f} = \frac{1}{Q_{10}}(14.9 + \frac{14.1}{1+exp\left(-\frac{Vm+95.2}{0.5}\right)}).$$

for the fast H current and

$$h_{s\infty}(V) = \frac{1}{1+exp(\frac{V+91}{10})}$$

and time constant

$$\tau_{h_s}(V) = \frac{1}{Q_{10}}(80 + \frac{172.7}{1+exp\left(\frac{Vm+59.9}{0.83}\right)})$$

for the slow H current.

**Passive currents (Leak and GABA-A)**

The model incorporated two different passive currents, the classical leak current, as introduced by Aradi and Holmes (Aradi and Holmes, 1999), and a tonic GABA-A chloride current, as introduced by Yim, Hanuschkin, and Wolfart in their model (Yim et al., 2015). The leak current represents a membrane's nonselective permeability to multiple ions and accounts for passive membrane properties in the Hodgkin-Huxley model. It was described as

$$I_L = \overline{g_L}(V - V_L)$$

Also, we adapted a published model incorporating the GABA-A chloride current (Yim et al., 2015) to account for the upregulation of the leak conductance in epileptic GCs (Young et al., 2009). It was described as

$$I_{GABAA} = \overline{g_{GABA-A}}(V - V_{GABAA}).$$

**Granule cell specific parameters**

The excitatory granule cell (GC), located in the granule cell layer, comprised a soma and a dendritic tree with two dendrites. Each dendrite was composed of 4 sections: the initial part of the dendrite located in the granule cell layer (GCLD), the proximal dendrite (PD), the medial dendrite (MD), and the distal dendrite (DD). The granule cell and its dendritic tree were depicted with 3D geometry and a location comparable to experimental data from a previous study (Tejada et al., 2014), which are available on the neuromorph site. To take into consideration the modification of intrinsic GC properties, parameters from a more recent study (Yim et al., 2015) were chosen. Passive parameters were:

| GC | Soma | GCLD | PD | MD | DD |
|---|---|---|---|---|---|
| Cm ($\mu F/cm^2$) | 0.75 | 0.75 | 1.2 | 1.2 | 1.2 |
| Ra ($\Omega$.cm) | 184 | 184 | 184 | 184 | 184 |

Reversal potentials were:

| GC<br>V (mV) | Soma, GCLD | PD | MD | DD |
|---|---|---|---|---|
| L | -83.8 | -81.74 | -81.74 | -81.74 |
| Na | 45 | 45 | 45 | 45 |
| K | -90 | -90 | -90 | -90 |
| GABA-A | -70 | -70 | -70 | -70 |

Maximal conductances were:

| GC<br>$\bar{g}$( $S/cm^2$) | Soma | GCLD | PD | MD | DD |
|---|---|---|---|---|---|
| NaT | 0.12 | 0.018 | 0.013 | 0.008 | 0 |

| GC | | | | | |
|---|---|---|---|---|---|
| NaP | $1.65 \cdot 10^{-5}$ | 0 | 0 | 0 | 0 |
| Kf | 0.016 | 0.004 | 0.004 | 0.001 | 0.001 |
| Ks | 0.006 | 0.006 | 0.006 | 0.006 | 0.008 |
| A | 0.012 | 0 | 0 | 0 | 0 |
| BK | 0.0006 | 0.0006 | 0.001 | 0.0024 | 0.0024 |
| SK | 0.001 | 0.0004 | 0.0002 | 0 | 0 |
| CaL | 0.005 | 0.0075 | 0.0075 | 0.0005 | 0 |
| CaN | 0.002 | 0.003 | 0.001 | 0.001 | 0.001 |
| CaT | $3.7 \cdot 10^{-5}$ | $7.5 \cdot 10^{-5}$ | $2.5 \cdot 10^{-5}$ | 0.0005 | 0.001 |
| L | $1.8 \cdot 10^{-5}$ | $1.8 \cdot 10^{-5}$ | $2.84 \cdot 10^{-5}$ | $2.84 \cdot 10^{-5}$ | $2.84 \cdot 10^{-5}$ |
| Kir | $7.22 \cdot 10^{-6}$ | $7.22 \cdot 10^{-6}$ | $7.22 \cdot 10^{-6}$ | $7.22 \cdot 10^{-6}$ | $7.22 \cdot 10^{-6}$ |
| GABA-A | $1.4 \cdot 10^{-5}$ | $1.4 \cdot 10^{-5}$ | $1.4 \cdot 10^{-5}$ | $1.4 \cdot 10^{-5}$ | $1.4 \cdot 10^{-5}$ |

**Mossy cell specific parameters**

The excitatory mossy cell (MC), located in the hilus, comprised a soma and a dendritic tree with four dendrites. Each dendrite was composed of 4 sections: the proximal dendrite (PD), the first part of the medial dendrite (MD1), the second part of the medial dendrite (MD2), and the distal dendrite (DD). Because of the low availability of 3D geometries of mossy cells in the literature, MCs were represented without any extent along the septotemporal axis. Passive parameters were:

| MC | | |
|---|---|---|
| | Soma | Dendrites |
| Cm (μF/cm$^2$) | 0 .6 | 2.4 |
| Ra (Ω.cm) | 100 | 100 |

Reversal potentials were:

| MC | |
|---|---|
| V (mV) | All compartiments |
| L | -59 |
| Na | 55 |
| K | -90 |
| Hf | -40 |
| Hs | -40 |

Maximal conductances were:

| MC | | | |
|---|---|---|---|
| $\bar{g}$( S/cm$^2$) | Soma | PD | others parts |
| NaT | 0.12 | 0.12 | 0 |
| Kf | $5{\cdot}10^{-4}$ | $5{\cdot}10^{-4}$ | 0 |
| A | $1{\cdot}10^{-5}$ | $1{\cdot}10^{-5}$ | $1{\cdot}10^{-5}$ |
| BK | 0.0165 | 0.00165 | 0.0165 |
| SK | 0.016 | 0.0016 | 0.016 |
| CaL | $6{\cdot}10^{-4}$ | $6{\cdot}10^{-4}$ | $6{\cdot}10^{-4}$ |
| CaN | $8{\cdot}10^{-5}$ | $8{\cdot}10^{-5}$ | $8{\cdot}10^{-5}$ |
| Hf | $5{\cdot}10^{-6}$ | $5{\cdot}10^{-6}$ | $5{\cdot}10^{-6}$ |
| Hs | $5{\cdot}10^{-6}$ | $5{\cdot}10^{-6}$ | $5{\cdot}10^{-6}$ |
| L | $1.3{\cdot}10^{-5}$ | $5.3{\cdot}10^{-5}$ | $5.3{\cdot}10^{-5}$ |

**Basket cell specific parameters**

The inhibitory basket cells (BC) comprised a soma and a dendritic tree with four dendrites: two apical dendrites and two basal dendrites. The apical dendrites were longer than the basal ones. Each type of dendrite was composed of the same sections: the proximal dendrite (PD), the first part of the medial dendrite (MD1), the second part of the medial dendrite (MD2), and the distal dendrite (DD). The basket cell and its dendritic tree were depicted with 3D geometry and a location comparable to experimental data from a previous study[13], available on the NeuroMorpho site. The soma of basket cells was located at the junction between the granule cell layer and the hilus. Passive parameters were:

| BC | |
|---|---|
| | All compartiments |
| Cm (μF/cm$^2$) | 1.4 |
| Ra (Ω.cm) | 100 |

Reversal potentials were:

| BC | |
|---|---|
| V (mV) | All compartiments |
| L | -60.06 |
| Na | 50 |
| K | -90 |

Maximal conductances were

| BC | | | | | |
|---|---|---|---|---|---|
| $\bar{g}$( S/cm$^2$) | Soma | PD | MD1 | MD2 | DD |
| NaT | 0.12 | 0.12 | 0 | 0 | 0 |
| Kf | 0.013 | 0.013 | 0 | 0 | 0 |
| A | $1.5{\cdot}10^{-4}$ | $1.5{\cdot}10^{-4}$ | $1.5{\cdot}10^{-4}$ | $1.5{\cdot}10^{-4}$ | $1.5{\cdot}10^{-4}$ |

| BC | | | | | |
|---|---|---|---|---|---|
| BK | 0.0002 | 0.0002 | 0.0002 | 0.0002 | 0.0002 |
| SK | $2 \cdot 10^{-6}$ | $2 \cdot 10^{-6}$ | $2 \cdot 10^{-6}$ | $2 \cdot 10^{-6}$ | $2 \cdot 10^{-6}$ |
| CaL | 0.005 | 0.005 | 0.005 | 0.005 | 0 |
| CaN | $8 \cdot 10^{-4}$ | $8 \cdot 10^{-4}$ | $8 \cdot 10^{-4}$ | $8 \cdot 10^{-4}$ | $8 \cdot 10^{-4}$ |
| L | $1.8 \cdot 10^{-4}$ | $1.8 \cdot 10^{-4}$ | $1.8 \cdot 10^{-4}$ | $1.8 \cdot 10^{-4}$ | $1.8 \cdot 10^{-4}$ |

**Hilar perforant path-associated cell specific parameters**

The hilar perforant path-associated cells (HC), located in the hilus, were inhibitory cells comprising a soma and a dendritic tree with 4 dendrites (2 shorts and 2 longs). Each dendrite was composed of 3 sections: the proximal dendrite (PD), the medial dendrite (MD), and the distal dendrite (DD). HIPP cells were represented without any extent along the septotemporal axis. Passive parameters were:

| HIPP | |
|---|---|
| | All compartiments |
| Cm (μF/cm$^2$) | 1.1 |
| Ra (Ω.cm) | 100 |

Reversal potentials were:

| HIPP | |
|---|---|
| V (mV) | All compartiments |
| L | -70.45 |
| Na | 55 |
| K | -90 |
| Hf | -40 |

| HIPP | |
|---|---|
| Hs | -40 |

Maximal conductances were:

| HIPP | | | |
|---|---|---|---|
| $\overline{g}$( $S/cm^2$) | Soma | PD | others parts |
| NaT | 0.2 | 0.2 | 0 |
| Kf | 0.06 | 0.06 | 0 |
| A | $8\cdot10^{-4}$ | $8\cdot10^{-4}$ | $8\cdot10^{-4}$ |
| BK | 0.003 | 0.003 | 0.003 |
| SK | 0.003 | 0.003 | 0.003 |
| CaL | 0.0015 | 0.0015 | 0.0015 |
| Hf | $1.5\cdot10^{-5}$ | $1.5\cdot10^{-5}$ | $1.5\cdot10^{-5}$ |
| Hs | $1.5\cdot10^{-5}$ | $1.5\cdot10^{-5}$ | $1.5\cdot10^{-5}$ |
| L | $4.7\cdot10^{-5}$ | $4.7\cdot10^{-5}$ | $4.7\cdot10^{-5}$ |

**Synaptic currents**

Excitatory and inhibitory synaptic currents followed

$$I_{syn} = \overline{g_{syn}} f(e^{-\frac{t+\Delta t}{\tau decay}} - e^{-\frac{t+\Delta t}{\tau rise}})\left(V - V_{syn}\right)$$

with $V_{syn}$ set to 0 mV for excitatory synapses and to -70 mV for inhibitory synapses, being the time after the arrival of a presynaptic spike and $\Delta t$ the synaptic delay,

$$f = e^{-\frac{tp}{\tau_{off}}} - e^{-\frac{tp}{\tau_{on}}}$$

being a normalization factor, with

$$tp = \frac{\tau_{off} * \tau_{on}}{\tau_{off} - \tau_{oon}} * log\left(\frac{\tau_{off}}{\tau_{on}}\right).$$

**Specific synaptic parameters**

Each synaptic type had specific values for maximal conductance, decay, rise, and delay. We also specified the synaptic location and the number of postsynaptic sites. In the following, excitatory synaptic currents were glutamatergic with a reversal potential $V_{syn} = 0$ mV, while inhibitory synaptic currents were GABAergic, with a reversal potential $V_{syn} = -70$ mV

**Perforant path synapses onto DG neurons**

In the model, perforant path (PP) fibers convey external spiking train inputs through excitatory synapses onto GC, MC, and BC neurons, with parameters set as :

| **PP-** | GC | MC | BC |
|---|---|---|---|
| $\overline{g_{syn}}$ (nS) | 20 | 0 | 10 |
| $\tau_{Rise}$(ms) | 1.5 | 1.5 | 2 |
| $\tau_{Decay}$(ms) | 5.5 | 5.5 | 6.3 |
| Delay (ms) | 3 | 3 | 3 |
| Synapse Location | DD | DD | DD |
| Number of postsynaptic sites | 2 | 4 | 2 |

**Synapses from mossy cells onto DG neurons**

Mossy cells projected excitatory synapses onto GC, MC, BC and HIPP neurons, with parameters set as :

| **MC-** | GC | MC | BC | HIPP |
|---|---|---|---|---|

| | | | | |
|---|---|---|---|---|
| $\overline{g}_{syn}$ (nS) | 0.3 | 0.5 | 0.3 | 0.2 |
| $\tau_{Rise}$(ms) | 1.5 | 0.45 | 0.9 | 0.9 |
| $\tau_{Decay}$(ms) | 5.5 | 2.2 | 3.6 | 3.6 |
| Delay (ms) | 3 | 2 | 3 | 3 |
| Synapse Location | PD | PD | PD Apicale | MD |
| Number of postsynaptic sites | 2 | 4 | 2 | 4 |

**Synapses from basket cells onto DG neurons**

Basket cells projected inhibitory synapses onto GC, MC and BC neurons, with parameters set as :

| **BC-** | | | |
|---|---|---|---|
| | GC | MC | BC |
| $\overline{g}_{syn}$ (nS) | 1.6 | 1.5 | 7.6 |
| $\tau_{Rise}$(ms) | 0.26 | 0.3 | 0.16 |
| $\tau_{Decay}$(ms) | 5.5 | 3.3 | 1.8 |
| Delay (ms) | 0.85 | 1.5 | 0.8 |
| Synapse Location | Soma | Soma | PD Apicale |
| Number of postsynaptic sites | 1 | 1 | 2 |

**Synapses from hilar perforant path-associated cells onto DG neurons**

Hilar perforant path-associated cells projected inhibitory synapses onto GC, MC and BC neurons, with parameters set as :

| HC- | | | |
|---|---|---|---|
| | GC | MC | BC |
| $\overline{g}_{syn}$ (nS) | 0.5 | 1.5 | 0.5 |
| $\tau_{Rise}$(ms) | 0.5 | 0.5 | 0.4 |
| $\tau_{Decay}$(ms) | 6 | 5 | 5.8 |
| Delay (ms) | 1.6 | 1 | 1.6 |
| Synapse Location | DD | MD | DD Apicale |
| Number of postsynaptic sites | 2 | 4 | 2 |

**Synapses from granule cells onto other DG neuronal sub-types**

Granule cells projected excitatory synapses onto GC, MC and BC neurons, with parameters set as :

| GC- | | | |
|---|---|---|---|
| | MC | BC | HIPP |
| $\overline{g}_{syn}$ (nS) | 0.2 | 4.7 | 0.5 |
| $\tau_{Rise}$(ms) | 0.5 | 0.3 | 0.3 |
| $\tau_{Decay}$(ms) | 6.2 | 0.6 | 0.6 |
| Delay (ms) | 1.5 | 0.8 | 1.5 |
| Synapse Location | PD | PD | PD |
| Number of postsynaptic sites | 4 | 4 | 4 |

**GC-GC (recurrent mossy fiber) synapses**

We adjusted GC-GC synaptic parameters for both AMPAR- and KAR-mediated synaptic responses to match experimentally recorded EPSPs and EPSCs (Artinian et al., 2011; Epsztein et al., 2010). Using a least-squares fitting method with $EPSC(V,t) = g_{syn}(t)(V_0 + EPSP(t))$, we obtained: $g_{AMPA} = 0.38\, nS$, $\tau_{Rise,\, AMPA} = 1.2\, ms$, $\tau_{Decay,\, AMPA} = 4.25\, ms$, $g_{KA} = 0.36\, nS$, $\tau_{Rise,\, KA} = 3\, ms$, $\tau_{Decay,\, KA} = 45\, ms$. In recurrent network simulations, synaptic maximal conductances were scaled to reproduce physiological AMPAR-EPSC amplitudes (30-35 pA)[14,15], while avoiding unrealistic connectivity (Santhakumar et al., 2005), $g_{AMPA} = 1.75\, nS$ (i.e., 4-5 times the one obtained by fitting experimental data). Based on experimental evidence that KAR-EPSCs are typically one-quarter to one-half the amplitude of AMPAR-EPSCs (Artinian et al., 2011; Epsztein et al., 2005; Matsuda et al., 2016; Pinheiro et al., 2013), we set $g_{KA} = g_{AMPA}/3 = 0.58\, nS$(Artinian et al., 2011; Epsztein et al., 2010, 2005).

| GC-GC | | |
|---|---|---|
| | AMPAR | KAR |
| $\overline{g_{syn}}$ (nS) | 1.75 | 0.58 |
| $\tau_{Rise}$(ms) | 1.2 | 3 |
| $\tau_{Decay}$(ms) | 4.25 | 45 |
| Delay (ms) | 0.8 | 0.8 |
| Synapse Location | PD | PD |
| Number of postsynaptic sites | 2 | 2 |

**Parameter estimation for NaP**

Previous work demonstrated that subthreshold KAR-EPSPs, but not AMPAR-EPSPs, activate persistent sodium currents (Artinian et al., 2011; Epsztein et al., 2010). We therefore incorporated a NaP current based on a published model (Lee, 2007). To simulate the maximal NaP amplitude in GCs, we used experimental data from (Epsztein et al., 2010). Using least-squares fitting between experimental curves and simulated currents described by $I_{NaP} = \overline{g_{NaP}} m_{\infty} h_{\infty} \left(V - V_{NaP}\right) S_{soma}$, where $S_{soma}$ is the somatic surface area, we determined the steady-state activation and inactivation parameters described (see NaP section above). To model activation kinetics, we introduced a scaling coefficient ($k_{NaP}$) to reproduce experimental activation time constants. This coefficient was estimated using a simplified GC model consisting of the sole soma with NaT, NaP, Kf, Ks, and leak currents. To compensate for the absence of dendritic potassium currents, Kf and Ks maximal conductances were increased by 25%. This simplified model exhibited a spike threshold of -55 mV and we performed measurements at the subthreshold holding potential of -60 mV. To simplify the kinetic parameter search, we initially assumed $\tau_{mNaP}$ to be voltage-independent and that the difference between resting and subthreshold EPSPs was entirely due to NaP. From recordings at resting potential, we estimated the passive parameters ($Cm$, $g_L$, and $V_L$) for each GC, using least-squares fitting of both AMPAR- and KAR-EPSPs at -60 mV. The resting condition corresponded to simulations without NaP, while the subthreshold condition included active NaP channels. We then optimized $g_{NaP}$ and $\tau_{mNaP}$ to minimize the difference between simulated and experimental AMPAR- and KAR-EPSPs simultaneously. This yielded $g_{NaP}$=0.155mScm$^{-2}$ and $\tau_{mNaP}$=40ms. We calculated $k_{NaP}$ such that $\tau_{mNaP}(V = -50mV) = 40ms$, giving $k_{NaP}$=525, allowing the model to reproduce experimental observations satisfactorily.

**Estimation of the local field potential (LFP)**

To assess whether the simulated activity was epileptiform, we computed an estimated extracellular local field potential (LFP) using the LFPy 2.0 toolkit. LFPy provides a Python interface to the NEURON simulation environment and includes dedicated methods to calculate extracellular potentials (Hagen et al., 2018; Hines et al., 2009). We therefore represented granule cells (GCs), mossy cells (MCs) and basket cells (BCs) with full 3D dendritic morphologies, as these geometries are required for reliable forward modeling of extracellular fields. The resulting simulated LFP signals were then compared with experimental LFP recordings to qualitatively assess the model's accuracy (Fig. 4).

**Neuronal and network observables**

All observables were computed based on granule cells' spike trains. The spike train of the k-th granule cell (over $n_{GC}$ ) was denoted $s_k$, with

$$s_k(t) = \sum_{l=1}^{n_k} \delta\left(t - t_k^l\right)$$

where δ is the Dirac delta function, $t_k^l$ the emission time of the $n_k$ spikes fired by the $k^{th}$ neuron, with spiking times $t_k^l \in \left[0, t_{max}\right]$, with $t_{max} = 310 + \Delta t_{window}(ms)$ being the time of simulation and $\Delta t_{window}$ the time window of stimulation of the network. The spike train of each neuron k was convolved by a Gaussian kernel with standard deviation 2.5 ms to get an estimate of the instantaneous firing frequency $f_k(t)$:

$$f_k(t) = s_k(t) \otimes \kappa(t)$$

with

$$\kappa(t) = \frac{1}{\sigma\sqrt{2\pi}} exp\left(-\frac{t^2}{2\sigma^2}\right).$$

The network mean spiking frequency was computed across neurons, i.e.,

$$\overline{f}(t) = \mu_k\left(f_k(t)\right),$$

where $\mu_x(.)$ is the mean over $x$. The maximal spiking frequency of the network was computed as

$$f_{max} = \max_t \left(\overline{f}_k(t)\right)$$

Sustained activity was assessed from $t_{start}$, i.e., which was defined as the first time at which the mean spiking frequency reached half the maximal spiking frequency

$$\overline{f}\left(t_{start}\right) = \frac{f_{max}}{2}.$$

In the following, we denote the initial and maximal times of the sustained period $t_0^{sust} \equiv t_{start}$ and $t_{max}^{sust} \equiv t_{max} - t_{start}$. Subsequent analysis was performed on sustained activity spike trains, i.e., between $t_0^{sust}$ and $t_{max}^{sust}$ and only for active neurons, i.e., who spiked more than 3 spikes during that period,

$$s_k^{sust}(t) = \sum_{l=1}^{n_k^{sust}} \delta\left(t - t_k^l\right)$$

with $t_k^l \in \left[t_0^{sust}, t_{max}^{sust}\right]$ the emission time of the $n_k^{sust}$ spikes fired by the $k^{th}$ of the $n_{GC}^{sust}$ active neurons during the sustained spiking period. In the following, all observables during that period are denoted with the sust superscript.

For each active neuron, we computed mean statistics over its inter-spike intervals (ISIs), i.e.,

$$ISI_k^l = t_k^{l+1} - t_k^l, \forall l \in \left[1, n_k^{sust} - 1\right], \forall k \in \left[1, n_{GC}^{sust}\right],$$

with $n_k^{sust}$ the number of spikes of the $k^{th}$ neuron. Specifically, we computed the mean spiking frequency of each individual neurons as

$$\overline{f_k^{sust}} = \frac{1}{n_k^{sust}-1}\sum_{l=1}^{n_k^{sust}-1} \frac{1}{ISI_k^l}$$

We also computed the CV2 of spiking activity, a local measures of spiking irregularity, for each neuron

$$CV_{2,k}^{sust} = \mu_l\left(2\frac{\left|ISI_k^l - ISI_k^{l-1}\right|}{ISI_k^l + ISI_k^{l-1}}\right)$$

The average mean of individual spiking frequency and irregularity were then computed as

$$\overline{f} = \mu_k\left(f_k^{sust}\right)$$

and

$$CV_2 = \mu_k\left(CV_{2,k}^{sust}\right).$$

The time constant of the autocorrelation of neurons' spiking activity was computed from spike lags in all neurons, defined as

$$lag_k^{i,j} = t_k^j - t_k^i, \forall i \in \left[1, n_k^{sust} - 1\right], \forall j \in \left[i + 1, n_k^{sust}\right], \forall k \in \left[1, n_{GC}^{sust}\right]$$

The probability distribution of time lags across all neurons was built as

$$p_{lag}^{t,t+\delta t} = \frac{Card_{\forall i,\forall j,\forall k}\left(t < lag_k^{i,j} \leqslant t + \delta t\right)}{Card_{\forall i,\forall j,\forall k}\left(lag_k^{i,j}\right)}$$

The time constant of the autocorrelation of spiking activity was obtained through an exponential fit of the probability distribution of time lags across all neurons during sustained spiking, for time lags from the peak of the probability distribution to its first null value. Neuronal spiking entropy was computed as

$$H_k = - \int p\left(f_k\right) log_2\left(p\left(f_k\right)\right) df_k,$$

$p\left(f_k\right)$ being the probability distribution of instantaneous frequency in neuron $k$ (obtained by convolution, see above). The average of neuronal entropy across neurons was then computed as

$$H = \mu_k\left(H_k\right).$$

Network synchrony was computed as

$$S = \sqrt{\frac{var_t\left(\mu_k\left(f_k^{sust}(t)\right)\right)}{\mu_t\left(var_k\left(f_k^{sust}(t)\right)\right)}}.$$

The dimensionality of network activity was computed as

$$D = \frac{\left(\sum_l \lambda_l\right)^2}{\sum_l \lambda_l^2}$$

where $\lambda_l$ are the eigenvalues of the normalized correlation matrix of instantaneous firing frequencies. Spiking mutual information between neurons $i$ and $j$ was computed as

$$MI_{i,j} = -\int_i \int_j p\left(f_i, f_j\right) log_2\left(\frac{p\left(f_i, f_j\right)}{p\left(f_i\right)p\left(f_j\right)}\right) df_i df_j,$$

$p\left(f_i, f_j\right)$ being the joint probability distribution of instantaneous frequencies in neurons $i$ and $j$. Network mutual information was then computed as

$$MI = \mu_i\left(\mu_{j>i}\left(MI_{i,j}\right)\right).$$

The cross-correlation space constant $\lambda$ was computed by fitting the dependence of the mean Pearson correlation coefficient between of GCs with other GCs, $C(d)$, as a function of their distance $d$ in the network:

$$C(d) = \frac{1}{n_{GC}} \sum_{i=1}^{n_{GC}} \frac{1}{2} (C_{i,i+d} + C_{i,i-d}),$$

with $d \in [1, 249]$ and

$$C_{ij} = \frac{cov_t(s_i(t), s_j(t))}{\sqrt{var_t\left(s_i(t)\right) var_t\left(s_j(t)\right)}},$$

by the exponentially decaying function

$$C_{fit}(d) = Aexp(-\frac{d}{\lambda}).$$

### Animal model of epilepsy and EEG recordings

Intra-hippocampal EEG recordings were obtained from a mouse model of temporal lobe epilepsy and digitized at 500 Hz. Epilepsy was induced in a seven-week-old male CD1 mouse (Charles River Laboratories) using the pilocarpine-induced status epilepticus model (Vigier et al., 2021). Following a chronic epileptic period (>2 months post-status epilepticus), the mouse was stereotaxically implanted with hippocampal electrodes as previously described (Peret et al., 2014). All experimental procedures were conducted in strict accordance with European Community Council Directive 2010/63/EU and approved by the French Ministry for Research following ethical review by the Aix-Marseille University Institutional Animal Care and Use Committee (protocol #9896).

### *References for methods*


Aradi I, Holmes WR. 1999. Role of multiple calcium and calcium-dependent conductances in regulation of hippocampal dentate granule cell excitability. Journal of Computational Neuroscience **6**:215–235. DOI: https://doi.org/10.1023/a:1008801821784, PMID: 10406134

Artinian J, Peret A, Marti G, Epsztein J, Crepel V. 2011. Synaptic Kainate Receptors in Interplay with INaP Shift the Sparse Firing of Dentate Granule Cells to a Sustained Rhythmic Mode in Temporal Lobe Epilepsy. Journal of Neuroscience **31**:10811–10818. DOI: https://doi.org/10.1523/JNEUROSCI.0388-11.2011

Epsztein J, Sola E, Represa A, Ben-Ari Y, Crépel V. 2010. A Selective Interplay between Aberrant EPSPKA and INaP Reduces Spike Timing Precision in Dentate Granule Cells of Epileptic Rats. Cerebral Cortex **20**:898–911. DOI: https://doi.org/10.1093/cercor/bhp156

Hagen E, Næss S, Ness TV, Einevoll GT. 2018. Multimodal Modeling of Neural Network Activity: Computing LFP, ECoG, EEG, and MEG Signals With LFPy 2.0. Frontiers in Neuroinformatics **12**:92. DOI: https://doi.org/10.3389/fninf.2018.00092, PMID: 30618697

Hines ML, Carnevale NT. 1997. The NEURON simulation environment. Neural Computation **9**:1179–1209. DOI: https://doi.org/10.1162/neco.1997.9.6.1179, PMID: 9248061

Hines ML, Davison AP, Muller E. 2009. NEURON and Python. Frontiers in Neuroinformatics **3**:1. DOI: https://doi.org/10.3389/neuro.11.001.2009, PMID: 19198661

Lee J. 2007. Fast Rhythmic Bursting Cells: The Horizontal Fiber System in the Cat's Primary Visual

Cortex.

Matsuda K, Budisantoso T, Mitakidis N, Sugaya Y, Miura E, Kakegawa W, Yamasaki M, Konno K, Uchigashima M, Abe M, Watanabe I, Kano M, Watanabe M, Sakimura K, Aricescu AR, Yuzaki M. 2016. Transsynaptic Modulation of Kainate Receptor Functions by C1q-like Proteins. Neuron **90**:752–767. DOI: https://doi.org/10.1016/j.neuron.2016.04.001

Molnár P, Nadler JV. 1999. Mossy fiber-granule cell synapses in the normal and epileptic rat dentate gyrus studied with minimal laser photostimulation. Journal of Neurophysiology **82**:1883–1894. DOI: https://doi.org/10.1152/jn.1999.82.4.1883, PMID: 10515977

Nörenberg A, Hu H, Vida I, Bartos M, Jonas P. 2010. Distinct nonuniform cable properties optimize rapid and efficient activation of fast-spiking GABAergic interneurons. Proceedings of the National Academy of Sciences of the United States of America **107**:894–899. DOI: https://doi.org/10.1073/pnas.0910716107, PMID: 20080772

Okazaki MM, Molnár P, Nadler JV. 1999. Recurrent Mossy Fiber Pathway in Rat Dentate Gyrus: Synaptic Currents Evoked in Presence and Absence of Seizure-Induced Growth. Journal of Neurophysiology **81**:1645–1660. DOI: https://doi.org/10.1152/jn.1999.81.4.1645

Peret A, Christie LA, Ouedraogo DW, Gorlewicz A, Epsztein J, Mulle C, Crépel V. 2014. Contribution of Aberrant GluK2-Containing Kainate Receptors to Chronic Seizures in Temporal Lobe Epilepsy. Cell Reports **8**:347–354. DOI: https://doi.org/10.1016/j.celrep.2014.06.032

Santhakumar V, Aradi I, Soltesz I. 2005. Role of mossy fiber sprouting and mossy cell loss in hyperexcitability: a network model of the dentate gyrus incorporating cell types and axonal topography. Journal of Neurophysiology **93**:437–453. DOI: https://doi.org/10.1152/jn.00777.2004, PMID: 15342722

Tejada J, Garcia-Cairasco N, Roque AC. 2014. Combined role of seizure-induced dendritic morphology alterations and spine loss in newborn granule cells with mossy fiber sprouting on the hyperexcitability of a computer model of the dentate gyrus. PLoS computational biology **10**:e1003601. DOI: https://doi.org/10.1371/journal.pcbi.1003601, PMID: 24811867

Vigier A, Partouche N, Michel FJ, Crépel V, Marissal T. 2021. Substantial outcome improvement using a refined pilocarpine mouse model of temporal lobe epilepsy. Neurobiology of Disease **161**:105547. DOI: https://doi.org/10.1016/j.nbd.2021.105547

Yim MY, Hanuschkin A, Wolfart J. 2015. Intrinsic rescaling of granule cells restores pattern separation ability of a dentate gyrus network model during epileptic hyperexcitability: INTRINSIC RESCALING AND NEURONAL NETWORK PATTERN SEPARATION. Hippocampus **25**:297–308. DOI: https://doi.org/10.1002/hipo.22373

Young CC, Stegen M, Bernard R, Müller M, Bischofberger J, Veh RW, Haas CA, Wolfart J. 2009.

Upregulation of inward rectifier K+ (Kir2) channels in dentate gyrus granule cells in temporal lobe epilepsy. The Journal of Physiology **587**:4213–4233. DOI: https://doi.org/10.1113/jphysiol.2009.170746, PMID: 19564397

***Supplementary figure 1***

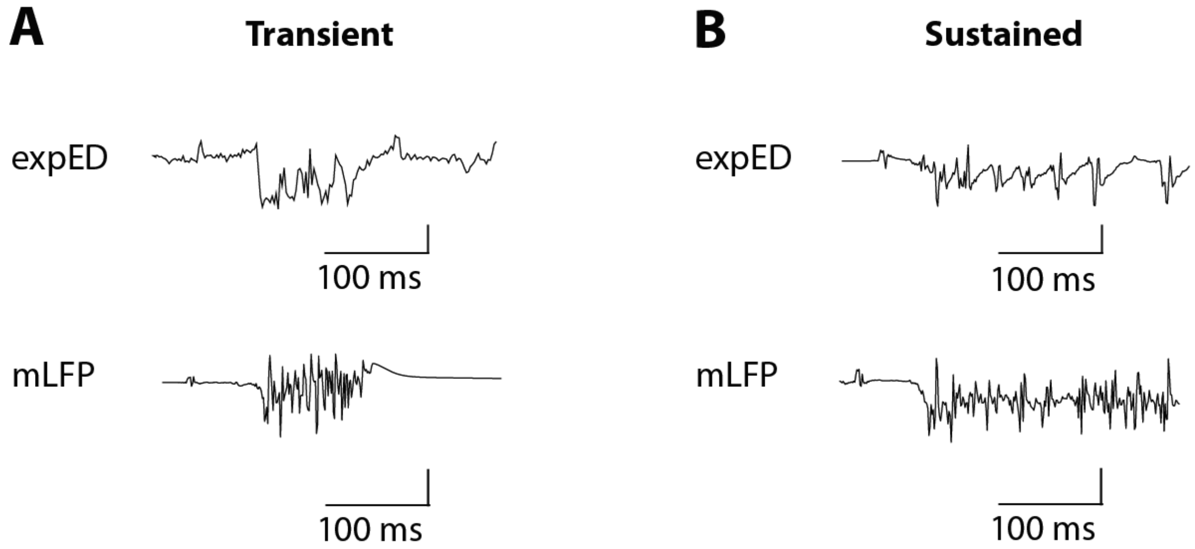


**Legend**

A and B show examples of experimental epileptiform discharges (expEDs) from a chronic epileptic mouse (top) and model local field potentials (mLFPs) (bottom). mLFPs are extracted from channels 1 and 9, as illustrated in Figures 4E and 4F, respectively. Note that mLFPs can display waveforms that resemble those of expEDs. Scale bars: 400 and 200 μV for expEDs and mLFP, respectively.
